# Learning within a sensory-motor circuit links action to expected outcome

**DOI:** 10.1101/2024.02.08.579532

**Authors:** WenXi Zhou, David M. Schneider

## Abstract

The cortex integrates sound- and movement-related signals to predict the acoustic consequences of behavior and detect violations from expectations. Although expectation- and prediction-related activity has been observed in the auditory cortex of humans, monkeys, and mice during vocal and non-vocal acoustic behaviors, the specific cortical circuitry required for forming memories, recalling expectations, and making predictions remains unknown. By combining closed-loop behavior, electrophysiological recordings, longitudinal pharmacology, and targeted optogenetic circuit activation, we identify a cortical locus for the emergence of expectation and error signals. Movement-related expectation signals and sound-related error signals emerge in parallel in the auditory cortex and are concentrated in largely distinct neurons, consistent with a compartmentalization of different prediction-related computations. On a trial-by-trial basis, expectation and error signals are correlated in auditory cortex, consistent with a local circuit implementation of an internal model. Silencing the auditory cortex during motor-sensory learning prevents the emergence of expectation signals and error signals, revealing the auditory cortex as a necessary node for learning to make predictions. Prediction-like signals can be experimentally induced in the auditory cortex, even in the absence of behavioral experience, by pairing optogenetic motor cortical activation with sound playback, indicating that cortical circuits are sufficient for movement-like predictive processing. Finally, motor-sensory experience realigns the manifold dimensions in which auditory cortical populations encode movement and sound, consistent with predictive processing. These findings show that prediction-related signals reshape auditory cortex dynamics during behavior and reveal a cortical locus for the emergence of expectation and error.

## Introduction

Anticipating how the world will change as we interact with it is foundational to behavior ^1–3^. Predictions allow animals to distinguish between environmental sensations (e.g. caused by other individuals) and self-generated sensations (i.e. caused by oneself) and are paramount to learning and executing behaviors such as speech and music ^4,5^. Theoretical frameworks for predictive processing propose the implementation of an internal model, which reflects a learned association between specific actions and their anticipated sensory outcomes; are built through experience; and are recalled during subsequent behavior to inform expectations and facilitate predictions ^6^. Behavioral and neural studies provide strong evidence for the use of internal models for sensory-motor processing in humans, monkeys, mice, fish, and other animals ^7–10^.

Building, storing, and recalling internal models that link action to outcome require integrating neural signals encoding behavior with neural signals encoding sensation. In hearing, movement signals interact with auditory signals throughout the neuroaxis, from the periphery to the cortex ^11^. As early as the middle ear, movement-related signals augment the gain of ascending auditory activity during behavior ^12,13^. But movement-related modulation becomes more nuanced in the auditory cortex, where it acts to selectively suppress neural responses to predictable self-generated sounds but not unexpected ones ^14–16^. Predictive signals are largely absent from the primary thalamus and in primary thalamo-recipient layers of the cortex (i.e. layer 4), suggesting that predictions arising from internal models may arise *de novo* in the cortex ^16,17^.

Within the auditory cortex, two prediction-like signals are consistent with the local implementation of an internal model. First, auditory cortical responses to expected self-generated sounds tend to be weak, whereas the same neurons’ responses to unexpected sounds tend to be large, consistent with the auditory cortex signaling a prediction error ^17–19^. Second, during a sound-generating behavior, movement-related signals in the auditory cortex become concentrated in neurons tuned to the expected sound and this activity peaks at the time the sound is expected to occur (even if the sound is omitted) ^16,17^. The auditory cortex thus encodes signals related to an animal’s experience-dependent expectation as well as whether the outcome of a behavior matched or violated the animal’s expectation (i.e. error).

Prediction-related signals in the auditory cortex are hypothesized to arise in part through its long-range, bidirectional connections with the secondary motor cortex (M2), a frontal region that is active during voluntary movements ^16,20–22^. Within the auditory cortex, M2 neurons synapse onto both excitatory and inhibitory neurons and many M2 neurons that innervate the auditory cortex have bifurcating axons that also target the brainstem via the pyramidal tract, thus allowing M2 to convey movement-related signals to auditory cortex via corollary discharge ^20^. Following extensive experience producing a sound-generating behavior, M2 dynamics change to reflect both movement and sound; and stimulation of M2 cells recruits distinct auditory cortex cells that are tuned to the sound a movement is expected to produce ^16,22^.

These studies provide strong but correlative evidence that the auditory cortex and its interactions with M2 play a role in learning, storing, and recalling internal models. However, it remains untested whether cortical circuits are necessary and sufficient for learning and recalling an internal model, nor is it known whether expectation- and error-like signals in the cortex interact on behavioral timescales needed to implement an internal model. Here, we combine closed-loop sound-generating behavior with dense, multi-time-point electrophysiological recordings and targeted circuit perturbations in mice learning the acoustic consequence of a novel behavior. We find that distinct prediction-related signals are carried by distinct auditory cortex populations, suggesting the compartmentalization of expectation and error signals within the auditory cortex. The emergence of a predictive phenotype requires the auditory cortex and artificial activation of motor cortical input to the auditory cortex is sufficient for learning prediction-like activity even in the absence of behavior. Once a prediction is formed, auditory cortex dynamics become reorganized such that movement signals align with dimensions that encode the expected consequences of action. These findings establish a cortical circuit that links an action to its expected outcome and reveal how movement, expectation, and error are encoded by single cells and populations of neurons in the cortex.

## Results

### Prediction-related signals emerge with motor-sensory experience

We trained head-restrained mice to make a simple and reproducible forelimb behavior (Fig. 1A). Mice held a lever with their right hand and pushed it away from a home threshold right in front of their body. Lever movements that passed a threshold located 12 mm from the home position were rewarded with a small drop of water when the mouse brought the lever back to the home position. All trials were self-paced and initiated by the mouse. Mice were initially trained on a silent lever with no auditory feedback, until their behavior reached an asymptote level of performance (Fig. 1B). We then modified the training configuration such that each lever movement produced a predictable sound (7kHz or 17kHz tone, randomized across mice) at a fixed position along the outgoing phase of the lever trajectory, without changing the reward contingency. We subsequently trained mice with the sound-generating lever for 7-8 days with 2 training sessions per day (Fig. 1B).

**Figure 1.**
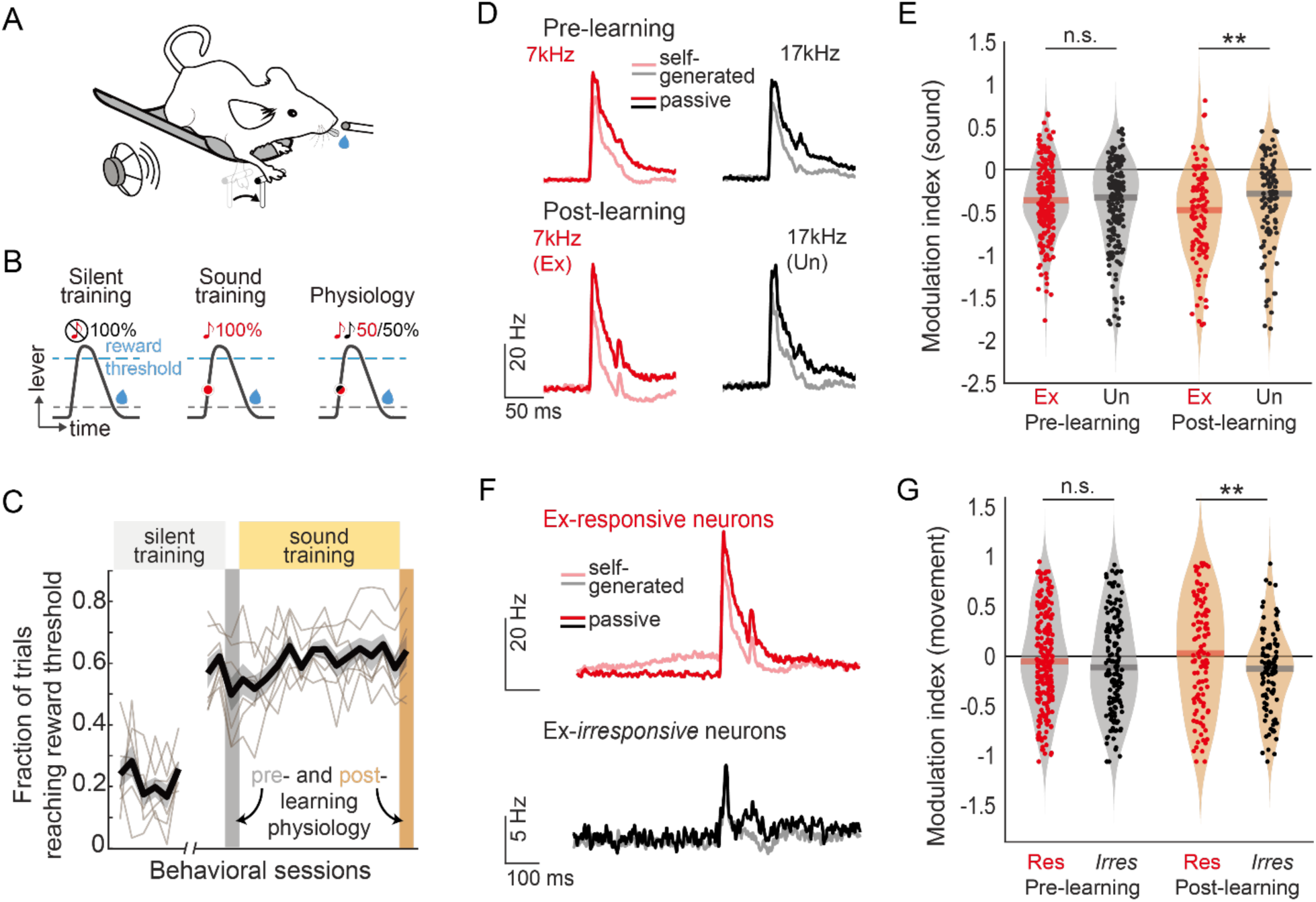
Prediction-related signals emerge with motor-sensory experience. **A.** Diagram of mouse operating a lever with its forelimb to receive a small water reward. Lever can either be silent or can be monitored in real-time to provide closed-loop auditory feedback. **B.** During silent training (left), lever movements that exceed a reward threshold (blue dashed line) are rewarded with a small drop of water when the mouse returns to the home position (black dashed line). During sound training (middle), a predictable sound (7kHz) is heard at a predictable position during every lever press. During physiology experiments (right), mice hear a sound at the predictable lever position, and the sound frequency is either 7 or 17kHz. **C.** Behavioral performance across training sessions. Physiological recordings were made just before and just after sound training. Behavioral performance was stable during physiological sessions and during sound training. **D.** Population-averaged neural responses to self-generated sounds before (top) and after (bottom) lever-sound learning. Dark lines show responses to self-generated tones, semi-transparent lines show responses to the same sounds heard passively. Responses to self-generated sounds are weaker than responses to passive sounds. After learning, responses to the expected sound (red) are markedly weaker than are responses to the unexpected sound (black). **E.** Quantification of suppression of self-generated tone responses before and after learning. Responses to both frequencies are equally suppressed before learning; after learning, responses to the expected sound are significantly more suppressed than responses to the unexpected sound. Red solid line indicates the mean suppression index. **F.** Population-averaged movement-related activity for neurons that are responsive to the lever-associated sound (red) and for neurons that are irresponsive to the lever-associated sound, but are responsive to other sounds (black). After learning, movement-related activity is concentrated in neurons that are responsive to the expected sound. **G.** Quantification of movement-related activity before and after learning. Before learning, movement-related activity is equivalent in neurons responsive to and irresponsive to the liver-associated tone. After learning, movement-related signals are significantly stronger in neurons responsive to the lever-associated tone.

We measured the responses of auditory cortical neurons to self-generated sounds with large-scale extracellular recordings at two distinct time points during behavioral training: at asymptotic performance but before introducing the lever-associated tone (pre-learning); and on the last day of behavioral training with the lever-associated tone (post-learning). During each of these recording sessions, mice pushed the lever and heard either a 7 or 17kHz tone with equal chance on every trial (Fig. 1B). At the end of each recording session, we also presented the same tones to the mouse during a passive listening period. Our experimental design provided two physiology sessions for each mouse during which mice had the same motor-sensory experience – pushing a lever and hearing one of two tones selected at random on each trial – with the important difference that during the post-learning session, mice expected the lever to produce one of the tones (expected) but not the other (unexpected). The mouse’s behavior remained stable throughout sound training and across both recording sessions (Fig. 1C).

Previous studies have identified two key features of auditory cortical activity that are consistent with a cortical role in predictive processing and that are present following motor-sensory experience ^17^. These physiological features include strongly suppressed responses to predictable self-generated sounds, which we refer to as predictive suppression, and movement-related signals that are concentrated in neurons tuned to the expected sound, which we refer to as movement-related expectation. We therefore first ensured that these signatures of predictive processing were absent in pre-learning recordings and only emerged following extensive motor-sensory training.

In pre-learning recordings, population-averaged neural responses to both of the self-generated tone frequencies were suppressed to a similar level compared to passive listening, consistent with the broadly reported finding of movement-related suppression of sound-evoked responses in the auditory cortex ^18,21,23^. In contrast, following extensive experience with the sound-generating lever, population-averaged responses became significantly more suppressed when the self-generated tone was expected, compared to when it was unexpected (Fig. 1D). We computed a single-cell modulation index and found that this population-level predictive suppression was recapitulated at the level of individual neurons (Fig. 1E). Predictive suppression was observed in both regular-spiking and fast-spiking neurons (which serve as signatures for putative excitatory and putative parvalbumin-positive interneurons, respectively) and was strongest in layer 2/3 compared to other cortical layers (Supp. Fig. 1).

We next asked whether and how movement-related signals in auditory cortex changed with experience. In both the pre- and post-learning recordings, nearly two-thirds of auditory cortical neurons were movement modulated – they changed their firing rates significantly just prior to or during the early phase of the lever movement (before the tone is heard) (Supp. Fig. 2A-D). Neurons with a fast-spiking waveform were more likely to be enhanced by movement, while regular-spiking neurons were equally likely to be enhanced or suppressed (Supp. Fig. 2E). In both pre- and post-learning recordings, neurons with strongly enhanced movement-related activity were more prominent in deep layers compared to superficial layers (Supp. Fig. 2F). In the pre-learning condition, movement-related signals were equally strong among neurons tuned to the lever-associated tone and in neurons tuned to other tones. In contrast, following experience with a sound-producing lever, the movement-related signals became concentrated in neurons that were tuned to the lever-associated tone and weaker in neurons tuned to other frequencies, reminiscent of a movement-triggered expectation (Fig. 1F-G).

### Expectation and error signals concentrate in distinct populations

Our two-time-point electrophysiology experiments reveal two hallmarks of predictive processing, the suppression of neural responses to predictable self-generated sounds and movement-related signals that reflect a feature-specific expectation. We next wondered whether these two predictive signals were concentrated in different neurons or whether the same auditory cortex cells encoded signals related to both expectation and error. To test this, we sorted neurons into two groups, those with significant movement-related changes in activity and those without. Neurons with strong movement-related signals did not tend to show predictive suppression - they had similar responses to both the expected and unexpected self-generated tones, and these responses did not change across the pre- and post-learning recordings (Supp. Fig. 3A). In contrast, predictive suppression was concentrated in neurons without movement-related activity and was only present after lever-sound training (Supp. Fig. 3B). This segregation of movement- and error-like signals into largely non-overlapping populations of neurons suggests a compartmentalization of functional roles during predictive processing in the cortex.

We next wondered whether these two prediction-related signals could be concentrated in even more refined subsets of neurons. Although our acute physiology recordings preclude us from following the same neurons across the pre- and post-learning sessions, we reasoned that functional clusters of auditory cortex cells could be identified and subsequently tracked across learning, providing an alternative approach to tracking the same neurons ^17,24^. To accomplish this, we took a data-driven approach to identify functional clusters of auditory cortex cells based solely on each neuron’s responses to passive sounds (see Methods). We combined non-negative matrix factorization (NMF) and k-means clustering to identify groups of neurons that could be consistently identified across mice and across both the pre- and post-learning recording sessions. We then analyzed the activity of these clusters during behavior to determine whether neurons with prediction-related signals (which can only be assessed during movement) can be identified by a neuron’s responses to passive sounds.

Our clustering strategy revealed 12 conserved groups of neurons with response profiles that varied along three principal dimensions: their time course of responses to passive tones (rapid decay versus slow decay), their relative tuning to expected and unexpected frequencies, and their direction of evoked responses (enhancement or suppression relative to baseline)(Supp. Fig. 5A). We found similar fractions of neurons belonging to each cluster before and after learning (Supp. Fig. 5B), consistent with previous findings that motor-sound experience does not alter neural tuning to passive sounds ^16^. Of these 12 clusters, we focused our analysis on 7 clusters that had rapid-onset, positive-going sound-evoked responses (Fig. 2A). The remaining clusters had negative-going sound-evoked responses or delayed responses that were inconsistent with classical sound processing, and their behavior-related activity was largely unaffected by auditory-motor experience (Supp. Fig. 5C).

**Figure 2.**
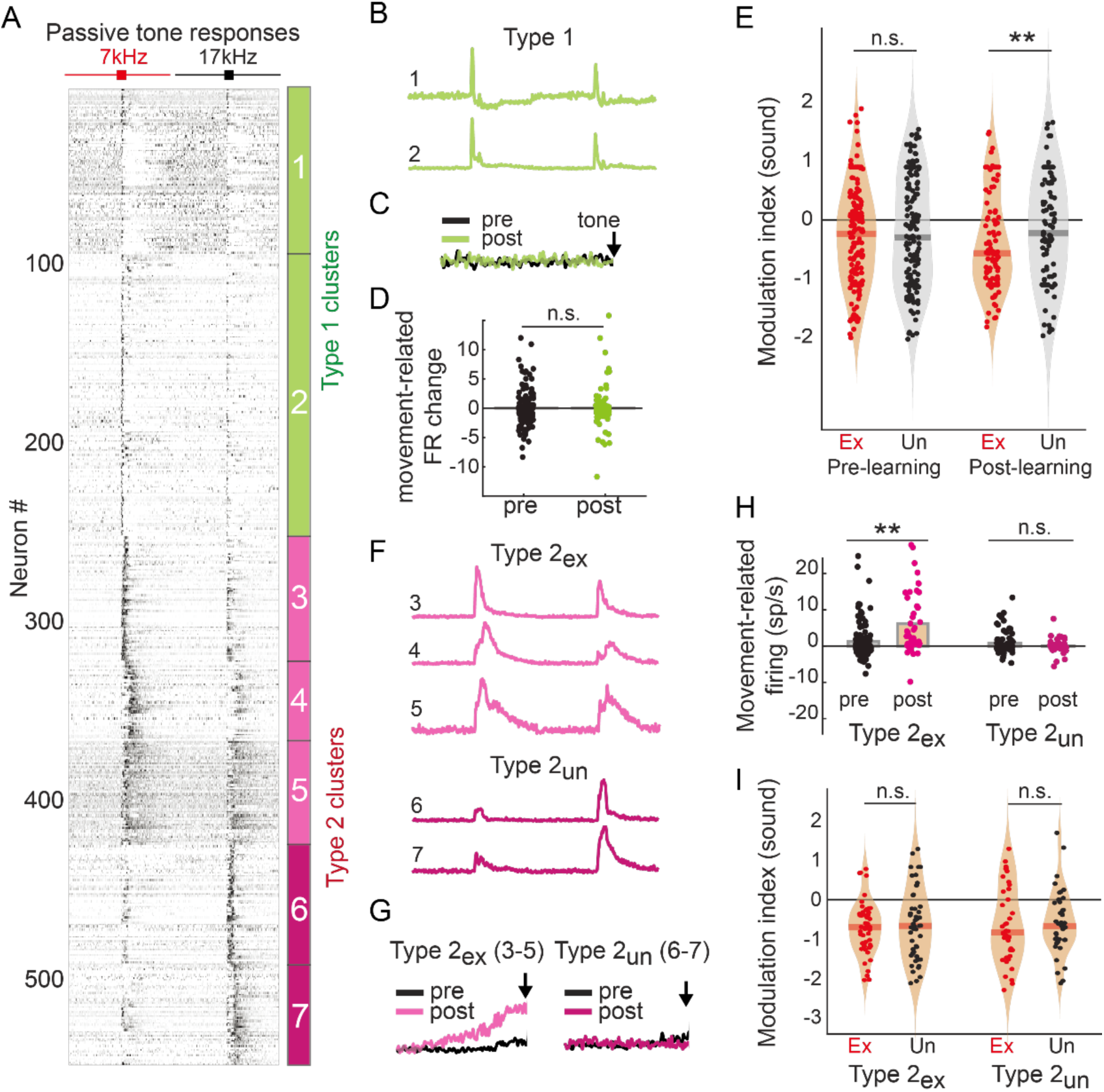
Expectation and error signals concentrate in distinct populations. **A.** Heat map showing concatenated neural responses to 7 and 17kHz tones heard passively. Neurons are organized into clusters using NMF and K-means. Data includes neurons recorded during both pre- and post-learning physiology. Only Type1 and Type2 clusters are shown. **B.** Population centroids for Type1 clusters (clusters 1 and 2). **C.** Population-averaged movement-related activity of Type1 clusters is minimal and is unaffected by lever-sound learning. **D.** Quantification of movement-related changes in firing rate from Type1 clusters during pre-learning (black) and post-learning (green) recordings. **E.** Type1 neurons develop frequency-specific suppression for the expected self-generated sound following lever-sound learning. **F.** Population centroids for Type2 clusters (clusters 3-7). Type2 neurons more strongly responsive to the lever-associated sound (3-5) and Type 2 neurons more strongly responsive to the unexpected sound (6-7) are further separated into Type 2ex (light pink) and Type 2un (dark pink). **G.** Population-averaged movement-related activity for Type 2ex and Type 2un neurons. After learning, Type 2ex but not Type 2un neurons have stronger movement-related activity. Black lines show movement-related activity before learning, pink lines show movement-related activity after learning. Black arrows indicate time of tone playback. **H.** Quantification of movement-related changes in firing rate from Type 2ex and Type 2un clusters during pre-learning (black) and post-learning (pink) recordings. **I.** Neither Type 2ex nor Type 2un neurons develop frequency-specific suppression for the expected self-generated sound following lever-sound learning.

The 7 sound-responding clusters that we identified could be further broken into two groups (Type1 and Type2) based on the time course of their passive sound responses. We identified two clusters that were characterized by rapid and brief tone-evoked responses (Type1) (Fig. 2B, Supp. Fig. 5D). These neurons were largely devoid of movement-related activity and exhibited predictive suppression of the expected self-generated sound during lever behavior (Fig. 2C-E). In contrast, Type2 neurons had rapid-onset but long-duration excitatory-like responses (Fig. 2F, Supp. Fig. 5D). These neurons tended to have positive movement-related activity that changed with lever-sound experience, but unlike Type1 neurons these cells did not exhibit predictive suppression (Fig. 2G-I). Both Type1 and Type2 units were dispersed across cortical depths, with no significant difference observed between them (Supp. Fig. 5E). These data indicate that aspects of a neuron’s sound-evoked response during passive listening, particularly the duration of evoked activity, is correlated with the presence/absence of movement-related activity and predictive suppression during behavior.

Type2 neurons could be further broken into 2 different groups, those that were more strongly responsive to the lever-associated tone (Type2-ex) and those that were more strongly responsive to the unexpected tone (Type2-un) (Fig. 2F). We predicted that experience with the tone-producing lever would lead to stronger movement-related responses in Type2-ex neurons, since these neurons received both movement- and sound-related input, which could potentially be involved in associative learning. Indeed, we found movement-related signals became stronger in Type2-exp clusters but not in Type2-un clusters (Fig. 2G-H). Meanwhile, sound responses of neither Type2-exp clusters nor Type2-un clusters changed with sensory-motor experience and showed relatively equal suppression towards expected and unexpected self-generated tones in both pre- and post-learning recordings (Fig. 2I). These clustering analyses reinforce our finding that movement-related expectation and predictive sound suppression are encoded by distinct populations of auditory cortex cells and identify an unbiased means of identifying neurons that encode prediction-related signals.

### Expectation- and error-related signals are correlated on behavioral timescales

Predictions about self-generated sounds are thought to arise through an internal model that compares expectation to experience on a trial-by-trial basis (Fig. 3A). Our data-driven clustering approach uncovered two distinct neuronal populations: one conveying expectation-like activity during movement and another representing an error-like signal following tone presentation. This result is consistent with local circuitry within the auditory cortex transforming movement-related input into an experience-related expectation that is used to modulate neural responses to upcoming self-generated sounds on a trial-by-trial basis (Fig. 3B). Based on this model, we predict that on trials when movement-related signals are more strongly concentrated in Type2-ex neurons the expectation will be strong, and thus tone-evoked responses (in Type1 neurons) will be weak.

**Figure 3.**
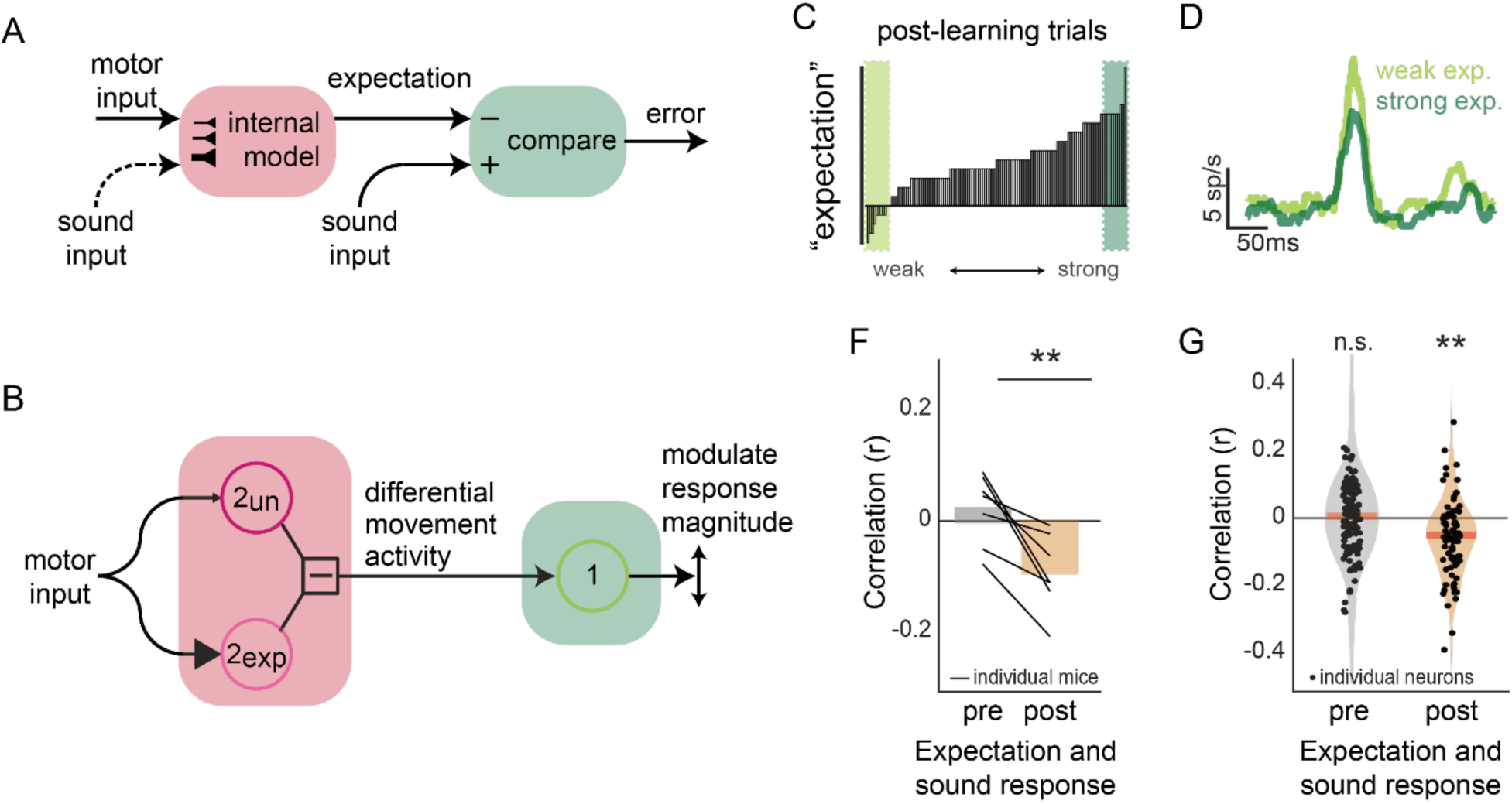
Expectation and error signals are correlated on behavioral timescales. **A.** A diagram for learning an internal model linking an action to its outcome, and for using expectation to compute errors. **B.** A diagram for Type 1 and Type 2 neurons in the auditory cortex could encode expectation. On a moment-by-moment basis, the difference in activity between Type 2ex and Type 2un neurons encodes an expectation, which in turn can modulate response magnitude of Type 1 neurons. **C.** Distribution of the difference in activity between Type 2ex and Type 2un neurons across all trials for an example mouse. Dark green shading shows trials with substantially higher activity in Type2ex neurons (i.e. strong expectation); light green shading shows trials with similar movement-related activity in Type 2ex and Type 2un neurons (i.e. weak expectation). **D.** Population-averaged response for Type 1 neurons on trials with weak and with strong expectations (see panel C). **E.** After learning but not before, on a trial-by-trial basis, gross movement-related activity (in Type 1 neurons) becomes anti-correlated with expectation-like activity (in Type 2 neurons). Each line shows an individual mouse, bars show average across mice. **F.** After learning but not before, on a trial-by-trial basis, expectation-like activity (in Type 2 neurons) becomes anti-correlated with the magnitude of evoked response to an expected self-generated sound (in Type 1 neurons). Each line shows an individual mouse, bars show average across mice. **G.** After learning but not before, on a trial-by-trial basis, the activity of individual Type 1 neurons is significantly anti-correlated with population-level expectation signals encoded by Type 2 neurons. Individual dots show individual Type 1 neurons.

To test this hypothesis, we performed a trial-by-trial analysis in which we compared the magnitude of the expectation signal on each trial with the magnitude of responses to the expected self-generated sound. To estimate expectation on each trial, we measured the difference between the magnitude of movement-related signals in Type2-ex and Type2-un populations: trials in which movement signals were similar in magnitude across both populations constituted a weak expectation, whereas trials in which movement signals were stronger in Type2-exp neurons constituted a stronger expectation. In post-learning sessions, movement signals were concentrated in Type2-exp neurons on most trials, consistent with an expectation for the lever-associated tone (Fig. 3C).

We next isolated trials in which the Type2 neural activity signaled a “weak” expectation and trials in which their activity signaled a “strong” expectation, and we analyzed sound-evoked activity in Type1 neurons independently in these two groups of trials. Consistent with our model, Type1 neural activity was weakest when Type2 neurons signaled a strong expectation and larger when Type2 neurons signaled a weak expectation (Fig. 3D). Across mice, we found that the magnitude of sound-evoked responses in Type1 populations was inversely correlated with the magnitude of movement-related expectation in Type2 populations, and that this relationship emerged only after motor-sensory training (Fig. 3E). In addition to these population-level correlations, we also analyzed single neuron correlations, comparing the sound-evoked activity of individual Type1 neurons to the strength of expectation encoded by simultaneously recorded Type2 populations (Fig. 3F). Even at the single-cell level, the magnitude of sound-evoked responses was significantly correlated with the magnitude of population-measured expectation. Together, these findings support a model in which movement-related signals reflecting expectation are used on a trial-by-trial basis to modulate neural responses to self-generated sounds and influence prediction-related activity.

### Auditory cortical activity is necessary for learning a predictive model

A prominent theory regarding the building of an internal model is that it is accomplished through activity-dependent changes in the strength of connections onto neurons that encode both action and outcome (see Fig. 3A). The presence of distinct populations of neurons in the auditory cortex that encode expectation and error suggests that plasticity required for building an internal model may occur locally within the auditory cortex. However, it remains untested whether the auditory cortex is necessary for learning the association between movement and its acoustic consequences.

To test this, we trained mice on the sound-generating lever while transiently silencing the auditory cortex during motor-sensory experience (sound training sessions), with topical, unilateral application of the GABA-A receptor agonist muscimol (see Methods) (Fig. 4A). We made physiological recordings from intact auditory cortex (without muscimol application) before and after training as in our standard experiments. The behavior of mice trained with their auditory cortex silenced remained stable across sound training (Fig. 4B). In post-learning sessions, auditory cortical responses to passive sounds were rapid and large, as observed in pre-learning recordings where no cortical silencing had ever been applied, suggesting that multi-day silencing of the auditory cortex did not alter cortical responses to passive sounds (Fig. 4C).

**Figure 4.**
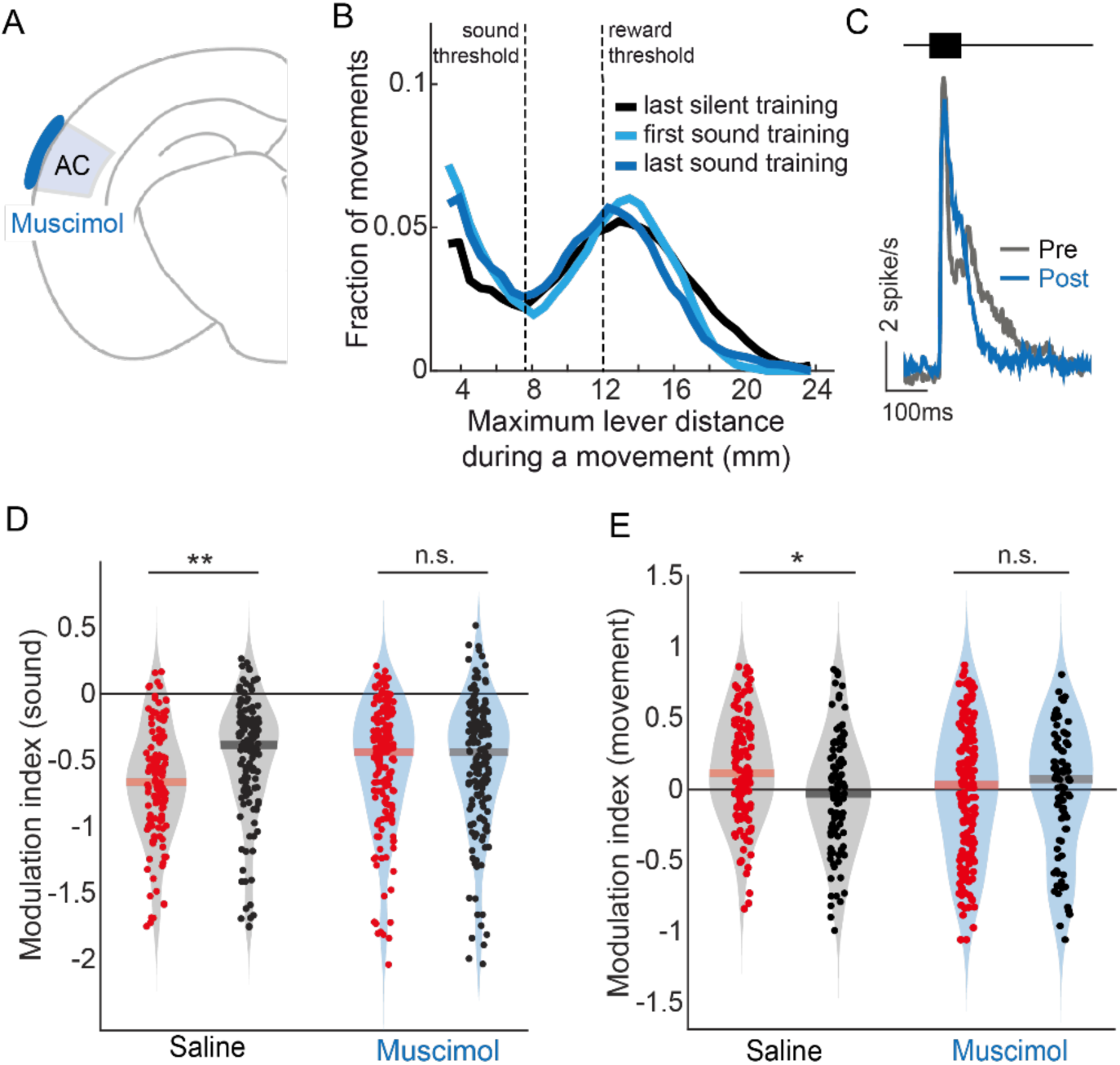
Auditory cortical activity is necessary for learning a predictive model. **A.** Diagram showing auditory cortical silencing during motor-auditory learning. **B.** Behavioral performance, measured as the peak lever position on each trial, is unaffected by muscimol application to auditory cortex. During silent training and during sound training with muscimol, the most common lever peak was at or just beyond the reward threshold. **C.** Longitudinal silencing of auditory cortex with muscimol does not impact subsequent auditory cortical responses to tones. Gray line shows neural responses before muscimol application (and before lever-tone training). Blue line shows neural responses after 7 days of muscimol application to auditory cortex during lever-tone training. **D.** Silencing auditory cortex during lever-sound training prevents the emergence of predictive suppression of expected tones in auditory cortex (blue). Control application of saline does not prevent the emergence of predictive suppression. **E.** Silencing auditory cortex during lever-sound training also prevents the emergence of expectation-like movement-related signals in the auditory cortex. Control application of saline does not prevent the emergence of expectation-like movement-related signals.

Silencing the auditory cortex during motor-sensory experience blocked the emergence of movement-related expectation and predictive suppression. In muscimol-trained mice, the suppression of sound-evoked responses during lever movements did not shift to become stronger for the expected lever-associated sound, while mice who received saline treatment instead of muscimol showed clear frequency-specific suppression, as what was observed in mice without any auditory cortex perturbation (Fig. 4D). Furthermore, with muscimol application during learning, movement-related activity prior to and early in the lever press remained uniform across auditory cortex cells, rather than concentrating in neurons tuned to the lever associated sound (Fig. 4E). The failure of these key neural signatures to arise indicates that auditory cortical activity during motor-sensory experience is necessary for the learning of an internal model that shapes activity at the single-cell and population levels to facilitate predictive processing.

### Fictive pairing of motor cortex and sound induces prediction-like activity in auditory cortex

After establishing that the auditory cortex is necessary for learning an internal model linking a specific action to its expected acoustic outcome, we next aimed to identify a long-range input that can provide movement-related signals and induce prediction-related learning in auditory cortex. We focused on the secondary motor cortex (M2), which sends long-range axons to the auditory cortex, synapses throughout cortical layers, drives strong suppression through feedforward inhibition, and has been implicated in predictive processing ^16,20,21^. M2 neurons, including those that synapse to auditory cortex are active during lever behaviors ^22^. However, it remains unresolved whether M2 is capable of driving the formation of an internal model and, once established, whether subsequent activation of M2 is sufficient to recall the internal model and drive prediction-like activity in auditory cortex.

We set to directly test this by pairing optogenetic M2 activation with tone playback (7kHz) when mice were not engaged in any task (Fig. 5A). On each trial, optical activation of M2 led passive tone presentation by 100ms, which mimics the delay between the lever initiation and self-generated tones in the sound-generating lever task. After 7-8 days (14-15 sessions) of pairing experience we recorded neural activity in auditory cortex while playing both the paired tone and tones of different frequencies, either with or without M2 activation. In general, optogenetic activation of M2 cell bodies led to both the activation and suppression of auditory cortical neurons within tens of milliseconds (Fig 5B), consistent with the pattern observed during the lever behavior and with a monosynaptic projection from M2 to auditory cortex ^20^. The proportion of downstream auditory cortical neurons significantly responding to optogenetic M2 activation is smaller than the proportion identified as movement-modulated during the sound-generating lever task (17% vs. 61%), which may be due to the spatial limitation of optogenetic stimulation or the lack of other coherent inputs received by auditory cortex during endogenous behavior (e.g. acetylcholine) ^25^.

**Figure 5.**
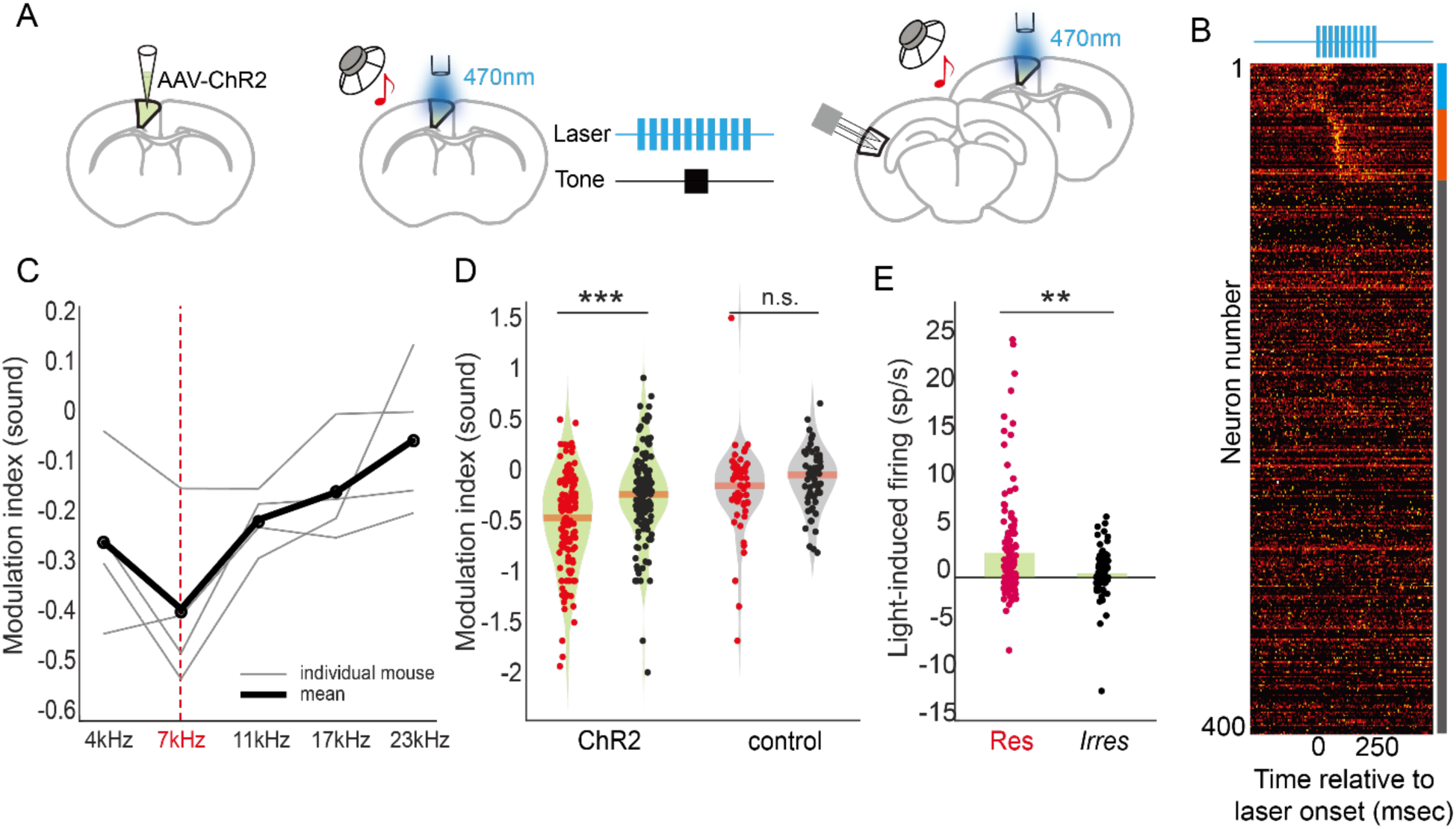
Fictive pairing of motor cortex activity and sound induces prediction-like activity in auditory cortex. **A.** Diagram showing infection of M2 neurons with ChR2 (left), pairing of M2 neurons with sound playback (middle), and physiological recordings from auditory cortex with sound playback and M2 stimulation. **B.** Heat map showing auditory cortical responses to optical M2 stimulation. Blue line on the right indicates neurons suppressed by M2 activation; red line indicates neurons enhanced by M2 activation; gray line indicates neurons that are unaffected. **C.** Following pairing of M2 activity and playback of 7kHz, subsequent M2 activation preferentially suppresses neural responses to 7kHz tones compared to other frequency tones, as indicated by notch in the modulation index at 7kHz. Gray lines show individual mice; black line shows average across mice. **D.** The modulation index of individual neurons shows specific suppression of 7kHz (paired) compared to 17kHz (unpaired); these sounds are used for direct comparison with mice who underwent lever-sound training. No frequency specificity arises following pairing of laser and sound in mice expressing an inactive fluorophore. **E.** Following pairing of M2 activation and 7kHz, subsequent M2 activation leads to a preferential recruitment of auditory cortical cells that are responsive to 7kHz, but not cells that are irresponsive to 7kHz (but are responsive to other tones).

When paired with sound-playback, M2 activation led to suppression of tone-evoked responses in most frequencies and importantly, a stronger suppression at the frequency that had been previously paired with M2 activation (Fig. 5C). As a more direct comparison with the experimental data collected during lever behavior, we compared neural responses to two frequencies – the 7kHz tone (paired) and 17kHz tone (unpaired) – in mice that experienced fictive pairing of M2 activation and sound playback. We found a signature of frequency-specific suppression with M2 activation that was qualitatively similar to the predictive suppression observed in lever-trained mice (Fig. 5D). Frequency-specific modulation by M2 activation was absent in control mice who underwent the same stimulation protocol but with an inactive fluorophore rather than ChR2 (Fig. 5D).

In addition to suppression of tone-evoked response, following repeated pairing of M2 activation with playback of a 7kHz tone, M2 activation more strongly recruited auditory cortical cells that were tuned to 7kHz (Fig. 5E). This biased distribution of the “movement-like” activity in auditory cortex after learning and the specific suppression of a sound frequency paired with M2 activation show that a circuit connecting M2 to auditory cortex is sufficient to build and recall an internal model linking action to acoustic outcome.

### Motor-sensory experience alters the geometry of cortical population dynamics

While movement-related activity is present throughout sensory cortices, population-level analyses have revealed that movement- and sensory-related signals tend to exist in dimensions that are largely orthogonal to one another, suggesting that behavior-related signals minimally impact the neural coding of sensation ^26–29^. However, prior experiments investigating the geometry of sensory cortical activity during behavior have largely focused on neural responses to sensations that are uncoupled to behavior. In our sound-generating lever task, actions consistently predict sensory feedback and experience reshapes how movement signals engage auditory cortical cells during behavior and sound processing. We thus wondered whether motor-sensory experience may alter the orthogonality between movement- and sensory-related dimensions of auditory cortex population activity.

To test this, we projected population-level activity from pre-learning recordings into a low-dimensional space using principal component analysis (Fig. 6A, Supp. Fig. 6A). Neural responses to sounds were concentrated in two principal components (PCs), with one dimension reflecting rapid-onset sound-evoked activity that did not distinguish among different sound frequencies (PC1) and the other reflecting sound-evoked activity with divergent patterns for each of the two frequencies studied here (PC3) (Supp. Fig. 6B). Movement-related signals were strongest in PC2, which showed weaker sound-evoked responses and was dominated by activity starting before and lasting throughout the lever movement (Supp. Fig. 6B).

**Figure 6.**
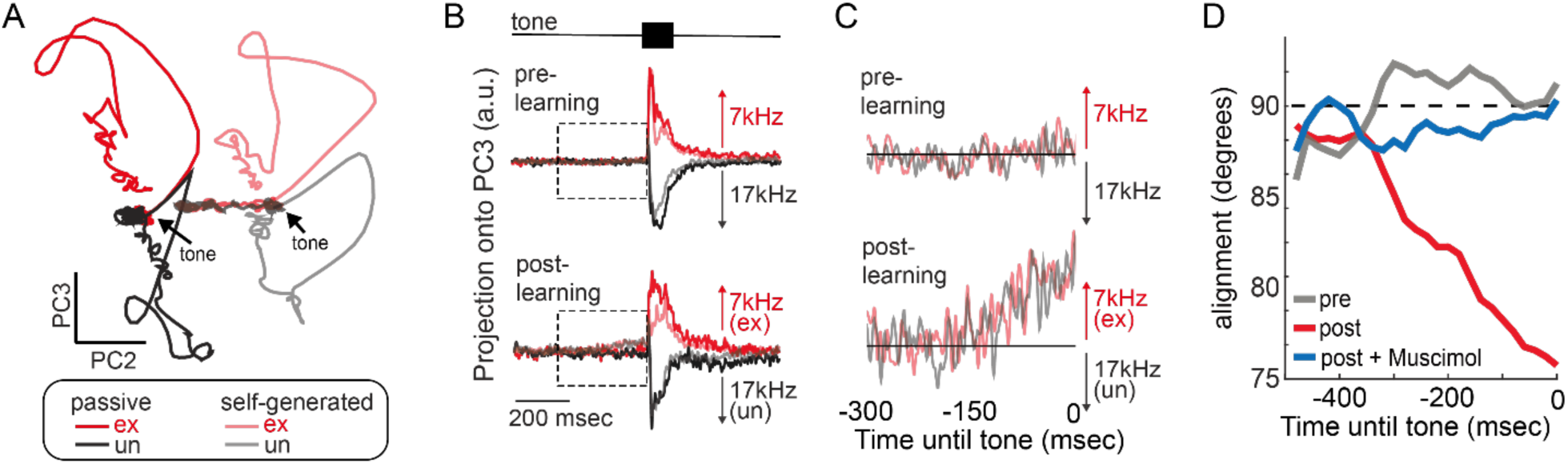
Motor-sensory experience alters the geometry of cortical population dynamics. **A.** Low-dimensional projection of auditory cortical population activity during passive sound playback of the lever-associated sound (7kHz, red) and another tone (17kHz, black). Light colors show neural responses to the same sounds when they are self-generated. Principal component 3 (PC3) distinguishes between the two sound frequencies whereas PC2 primarily encodes movement. Data are from electrophysiological recordings made before the lever was expected to produce 7kHz (pre-learning). **B.** Projection of neural activity along PC3 from pre-learning (top) and post-learning (bottom) recordings. At the time of the tone, PC3 distinguishes between the two sound frequencies. Dashed box indicates zoomed-in time window in panel C. **C.** Zoom in to dashed box in panel B showing activity in PC3 before the tone is heard (time 0). Before learning (top), movement-related activity is largely absent from PC3. After learning (bottom), movement-activity is present in PC3, and it specifically pushes population dynamics toward the dimension encoding the sound the lever is expected to produce. **D.** The angle between ongoing population dynamics before sound presentation (i.e. movement-related activity) and the dimension that separates expected frequency from unexpected frequency during passive listening (frequency dimension). Before lever-sound training (pre, gray), movement-related activity is orthogonal to the frequency dimension. Following lever-sound training (post, red), movement-related activity becomes more aligned with frequency dimension. Application of muscimol to auditory cortex during lever-sound training prevents the experience-dependent realignment of movement and sound dimensions (post + Muscimol, blue).

In pre-learning recordings, movement-related signals were present in PC1, which reflects frequency-independent sound-evoked activity, but were largely absent from PC3, which encodes sound frequency (Fig. 6B-C, top panel). These findings are consistent with movement acting grossly on sensory-encoding dimensions that are feature-independent, but not interacting with sensory dimensions that encode specific sensory features^26^. In contrast, following learning we observed movement-related activity in PC1 as well as in PC3 (Fig. 6B-C, bottom panel). Importantly, the movement-related activity in PC3 specifically pushed neural dynamics in the direction of the sound frequency the lever was expected to produce, consistent with movement signals encoding an expectation for the lever-associated tone (Fig. 6C).

To quantify this more thoroughly, we computed the angle between the trajectory of movement-related signals and the dimension that best separates the sound frequencies using the full dimensional space rather than just the first three PCs. In pre-learning recordings, the movement and sound-frequency dimensions remained orthogonal to one another (Fig. 6D). In contrast to the pre-learning recordings, following learning we observed a realignment of movement- and frequency-encoding dimensions such that neural trajectories during movement (but prior to the self-generated tone) were no longer orthogonal to the plane that separates the expected and unexpected frequencies. Instead, this realignment biased movement-related activity in the direction representing the sound frequency associated with the lever, revealing an expectation-like signal in the population dynamics of auditory cortical activity (Fig. 6D).

As a final analysis, we examined population dynamics in mice that had their auditory cortex silenced with muscimol during motor-sensory experience. In these mice, the movement- and frequency-dimensions remained largely orthogonal to one another, despite substantial motor-auditory experience (Fig. 6D). Together, these findings reveal that with experience, movement-related signals in auditory cortex take on an expectation-like phenotype that may reflect a cortically based internal model. The building and recall of such an internal model requires auditory cortical activity during experience.

## Discussion

Using a simple closed-loop behavior, we find that predictive processing in the auditory cortex involves learning within a distributed sensory-motor cortical circuit. We find that experience producing a movement with a predictable acoustic outcome leads to two prominent prediction-related signals in auditory cortex: expectation-like movement-related activity and frequency-specific suppression of tone-evoked responses. While both signals have been observed previously ^17^, by making physiological recordings from the same mice before and after learning, we show that these signals only emerge following extensive motor-auditory experience. We further find that different populations of neurons encode signals related to expectation and error and that activity of these populations is correlated on a trial-by-trial basis. The emergence of prediction-related signals in auditory cortex requires auditory cortical activity during motor-sensory experience and could be recapitulated by pairing M2 activation with sound playback. Finally, we find that motor-sensory experience reshapes population dynamics in the auditory cortex by aligning movement-encoding and sound frequency-encoding dimensions.

### Expectation- and error-related signals are concentrated in distinct groups of neurons

Using a combination of dimensionality reduction (NMF) and clustering (k-means) techniques, we found that two prediction-related signals were concentrated in different clusters of neurons in auditory cortex and that these clusters could be identified by a neuron’s responses to passively heard tones. This segregation of signals provides multiple insights regarding the mechanisms of predictive processing in the cortex.

First, the segregation of expectation-like movement-related activity and suppressed tone-evoked responses ruled out a mathematically trivial relationship between these two signals. In particular, we measure tone-evoked responses relative to a neuron’s activity just before tone onset. If this activity was elevated for neurons tuned to the lever-associated tone, this could lead to weaker tone-evoked responses simply because the two phenomena are strongly coupled for individual neurons. In contrast, the decoupling of these signals throughout the population reveals that different auditory cortical cells encode two distinct prediction-related signals during behavior. These two prediction-related signals likely represent consecutive stages of predictive processing. Movement-related input is first received by one group of neurons that shows activity changes early in behavior, and the output of these cells encodes an expectation-like signal that is used to suppress neural responses to the expected tone in a separate group of error-detecting neurons. Indeed, on a trial-by-trial basis we found that the concentration of movement-related activity in neurons tuned to the expected frequency was correlated with the strength of suppression to the expected self-generated sound. This trial-by-trial correlation is consistent with the implementation of an internal model within the auditory cortex on rapid, behavioral timescales.

Second, it is notable that different populations of neurons that encode expectation-like and error-like signals – both of which can only be measured during behavior – could be identified using only sound responses during passive listening. This finding suggests that there may be basic functional groups of neurons in auditory cortex that are recruited during predictive processing, complementing recent work showing that prediction-related neurons in visual cortex have distinct molecular profiles ^30^. Here, we found that movement-related expectation-like signals are concentrated in neurons with longer-duration sound-evoked responses. Strong and prolonged sound-evoked activity may permit a longer window during which transient and potentially weak presynaptic motor input could coincide with postsynaptic sound responses. Our data are consistent with a potential mechanism for building an internal model through experience-dependent changes in the efficacy of motor-related inputs onto auditory cortical cells that exhibit prolonged responses to self-generated sounds.

In future experiments, it will be informative to monitor the activity of the same neurons longitudinally (e.g. using 2-photon calcium imaging) as mice learn to predict the acoustic consequences of a movement. By following the same neurons across days, such experiments can help clarify whether auditory cortical cells move from one functional cluster to another during learning, or whether cluster identity stays constant while the functional signals encoded by those clusters change over time. It also remains unknown whether the functionally identified prediction-related clusters of neurons we identified here correspond to the molecularly identified clusters of neurons that have previously been implicated in predictive processing ^30^. Finally, the circuit logic for comparing expectation with experience is currently better understood in other neural systems, such as the cancellation of self-generated electrical input in weakly electric fish ^31,32^. It will be important to understand the local circuit logic within the auditory cortex that compares movement-related activity across two groups of neurons and uses the output of this comparison to modulate sound-evoked responses in a separate population.

### Silencing auditory cortex prevents the building and recall of an internal model

Movement and expectation influence neural processing across many stages of the auditory pathway, from the ear to the cerebral cortex ^11,33^. For example, sound-evoked responses are suppressed in the auditory thalamus during behavior ^16^, movement-related signals are present in the inferior colliculus ^34,35^, and self-generated sounds are suppressed as early as the dorsal cochlear nucleus (DCN) ^13^. The presence of movement-related signals along the auditory neuroaxis has raised the question of where along this pathway prediction-related signals arise. By silencing the auditory cortex during motor-sensory experience, we show that prediction-related signals arise in the auditory cortex and that auditory cortical activity is necessary for the emergence of predictive signals. This is the first causal manipulation to show that mouse auditory cortex is necessary for the emergence of prediction-related signals and it is consistent with previous studies reporting an absence of such signals in the auditory thalamus ^16^ or L4 of primary auditory cortex ^17^, and the necessity of NMDA-receptors in the mouse visual cortex for visuomotor integration ^36^.

Given that the predictive suppression in auditory cortex is found along multiple dimensions including frequency, timing, intensity, and spatial location ^19^, it is perhaps not surprising that the auditory cortex – which encodes all of these signals – is required in the formation of internal model. Previous work has found that prediction error signals in mouse auditory cortex occur coincidentally with sound responses, ruling out feedback from downstream areas as a source of prediction error signals in auditory cortex ^19^, further pointing to the auditory cortex as a site of learning an internal model. Yet the experiments performed here cannot dismiss the possibility that plasticity required for forming an internal model may occur outside of the auditory cortex (e.g. in downstream areas), and experiments that preserve auditory cortical activity but block plasticity will be needed to definitively pinpoint the locus of plasticity.

### M2 activation is sufficient to build and drive prediction-like signals in auditory cortex

Previous studies have suggested that M2 could play an important role in the predictive processing of self-generated sounds. M2 neurons send long-range axons to the auditory cortex, where they make excitatory synapses onto inhibitory and excitatory cells across most cortical layers ^20^. Artificially activating M2 axons is sufficient to suppress auditory cortical membrane potential activity in a manner similar to locomotion and turning M2 off during locomotion reverts auditory cortex activity to a rest-like state ^21^. And following motor-sensory experience, stimulating M2 results in the preferential recruitment of auditory cortical cells tuned to the movement-associated sound, providing correlative evidence for M2 in predictive processing ^16^. However, prior studies have not used causal perturbations to test whether M2 is sufficient for learning and subsequently driving prediction-like signals in auditory cortex.

Our fictive pairing experiment shows that M2 activation alone can induce the formation of an internal model. The expectation- and error-like signals that emerge following fictive pairing are qualitatively and quantitatively similar to the prediction-related signals that emerge following normal lever-sound experience. While more nuanced experiments will help pinpoint the specific connections from M2 to auditory cortex that drive learning (e.g. axon terminal manipulations), our work identified M2 activity as being capable of driving prediction-like signals even in the absence of behavior. In future work, it will be important to understand the roles that other behavior-related inputs play in the formation and recall of internal models, such as the cholinergic inputs to auditory cortex that are active during locomotion and other behaviors ^25^.

### Predictive processing through the lens of neural dynamics

We used population dynamics as a means of visualizing and understanding how motor- and sound-related signals interact in the auditory cortex before and after mice form an expectation. Movement signals have been observed throughout the mouse brain, particularly in sensory and associative regions of the cortex ^21,37–39^. Consistent with previous reports of behavior-related signals in sensory cortex, we found that before lever-sound learning, movement-related signals in auditory cortex are primarily orthogonal to activity dimensions that encode stimulus features (e.g. sound frequency) ^26^. However, following motor-sensory experience, movement-related signals become more aligned with the activity dimension that encodes stimulus frequency. In particular, movement signals move in the direction that encodes the expected frequency but not the unexpected frequency. This alignment of activity dimensions encoding movement and acoustic features shows that auditory cortical dynamics can be shaped by experience to reflect motor-sensory expectations.

The prevalence of movement-related signals in sensory areas even without a motor-sensory association, and the reshaping of these signals when an association emerges, suggests that movement-related signals might serve multiple roles in modulating sensation and perception during behavior. In situations where sensory input is independent from an animal’s behavior, spontaneous and movement-related dynamics push population activity away from basal states without intruding upon sensory-encoding dimensions, which could serve as a context cue for discriminating the conditions in which stimuli are observed ^40^. Our single-neuron and population-level analyses suggest that during behaviors with sensory outcomes, movement-related signals could provide a teaching signal for learning the expected outcome of a particular action. Finally, once a stable association between action and outcome has been established, movement-related signals become aligned with specific sensory features, which could allow easier processing of expected sensory inputs and facilitate detection of deviant outcomes.

## Acknowledgements

We thank Grant Zempolich and Ariadna Corredera for their feedback on this manuscript and members of the Schneider lab for their input throughout the project. We thank SueYeon Chung and Aishwarya Balwani for conversations regarding population coding and neural manifolds. We thank Hoda Ansari, Jessica A. Guevara, and Deanna Garcia for their assistance with managerial support and animal care. This research was supported by the National Institutes of Health (1R01-DC018802 to D.M.S.); a Career Award at the Scientific Interface from the Burroughs Welcome Fund (D.M.S); fellowships from the Searle Scholars Program, the Alfred P. Sloan Foundation, and the McKnight Foundation (D.M.S.). D.M.S. is a New York Stem Cell Foundation - Robertson Neuroscience Investigator.

## Methods

### Experimental model and subject details

All experimental protocols were approved by New York University’s Animal Use and Welfare Committee. Wild-type (C57BL/6) mice were used for all other experiments. Mice were purchased from Jackson Laboratories and subsequently bred in an onsite vivarium. We used 2–4 months old male and female mice for our experiments. Mice used for experiments were kept on a reverse day-night cycle (12h day, 12h night).

### Surgeries

For all surgical procedures, mice were anesthetized under isoflurane (1-1.5% in O2) and placed in a stereotaxic holder (Kopf). Both the skin over the top of the head and the muscle over the left auditory cortex were removed during surgery. The left auditory cortex (AP: -2.8mm, ML: -4.4mm) was marked on the skull with black ink for a subsequent craniotomy. For all mice, a Y-shaped titanium headpost (H.E. Parmer) was attached to the skull using a transparent adhesive (Metabond). Meloxicam SR was applied subcutaneously post-surgery as an analgesic. All mice were allowed to recover for at least 3 days prior to the start of water restriction. Following training and 12 to 24 hours prior to the first electrophysiology recording, a small craniotomy (∼2mm diameter) was made over the auditory cortex according to the marker made on the skull. Exposed craniotomies were covered with a silicone elastomer (Kwik-Sil). Another small craniotomy was made above the right sensory cortex and a silver-chloride reference electrode was positioned atop the surface of the brain for use as a ground electrode and covered with dental cement (AM Systems).

For mice used for fictive pairing experiments, a small craniotomy was made over the left M2 (AP: 1.4mm, ML: -0.8mm) during the headpost implanting surgery. A total of 150nL virus expressing ChR2 (CaMKIIa-hChR2-mCherry, UNC) or mCherry (CaMKIIa-mCherry, Addgene114469) was injected at three depths (0.3, 0.6 and 0.9mm below dura) using an injector (Drummond Nanoject) with a pulled glass pipette (Drummond). A custom-made optic fiber assembly (Thorlabs, FP400URT and SFLC440) was then implanted at the same coordinates right above the surface of the brain and was secured using Metabond together with the headpost. Mice were allowed 3 weeks prior to the start of pairing for recovery and viral expression. Mice used for fictive pairing experiments were not water restricted.

### Sound-generating lever task

We used a head-restrained lever-based behavioral training paradigm where mice push a lever and hear closed-loop sounds. The lever rig and the lick port used to deliver water rewards were custom designed (see Audette et al., 2022). Digital signals for lever movement were collected by a data acquisition card (National Instruments) connected to a computer, logged by custom Matlab software (Mathworks, PsychToolBox) and used to trigger reward and sound events. Sound output was delivered from an ultrasonic speaker (Tucker Davis Technologies) positioned close to the mouse’s right ear through a sound card (RME Fireface UCX). All training was performed in a sound-attenuating booth (Gretch-Ken) to minimize background sound and monitored in real-time via IR video.

During the lever task, mice were water restricted, maintained at greater than 85% of pre-restriction body weight, and received all of their water (1-2mL per day) while performing the lever behavior. Mice were first trained with a silent lever for approximately three weeks with two sessions per day to acquire a stereotypical lever pushing behavior. At the beginning of silent training, the reward threshold was set very close to the starting position and mice received a water reward (∼5uL) for each lever press that crossed the reward threshold; rewards were delivered when the lever was brought back to the starting position. The reward threshold was gradually moved away from the mouse’s body to a final position ∼12mm away from the starting position, which was achieved after around 1 week of training. The reward threshold then stayed the same for the rest of training and recording. During silent training, the maximum reach of each lever press was monitored and the distribution of the maximum reach was used as an indicator of the mouse’s task performance. During the later stages of silent lever training, mice were also trained to obey an inter-trial interval (ITI) such that only trials initiated 500ms after the end of the previous trial were rewarded. Mice were also not allowed to lick during the ITI or the reward was omitted. We considered mice qualified for electrophysiology and sound-lever training when they satisfied two criteria. First, mice needed to receive at least 150 rewarded trials during the ∼1 hour session, indicating that their behavior obeyed the ITI rules. Second, we required that the distribution of peak lever-press distances formed a Gaussian-like distribution with a peak around the reward threshold that was stable across days. Once mice met these requirements, a 50ms pure tone (7kHz or 17kHz, randomly chosen for each mouse) was introduced and paired with lever presses at ∼6mm away from the starting position (sound threshold). The sound threshold was deliberately chosen to be closer to the body than the reward threshold to maintain a high probability of hearing the tone given a lever press behavior. For each lever press that reached further than the sound threshold, the chosen pure tone would be played immediately after the threshold crossing. All other behavioral criteria were maintained throughout sound training. Mice experienced 14-15 sessions of sound training over one week.

Right before and after the introduction of sound training, we assessed the mouse’s expectation for self-generated sounds and recorded neural activities in auditory cortex. For the first 5-10 minutes of the recording session, mice had the same experience as sound training, with each lever push triggering only the chosen pure tone. Trials from this habituation period of time were not used for later analysis. Then a probing tone with a different frequency (7kHz or 17kHz, depending on the previously chosen tone) was introduced and triggered by 50% of lever presses instead. The mice were allowed to move the lever by will during the whole recording session; but in post-hoc, any trial that didn’t obey the ITI rule (no lever movement and licking 500ms prior to trial initiation) was excluded. In addition, in post-hoc, if the mice triggered a second tone within 200ms from the offset of the previous tone, the previous trial was excluded. This inclusion criteria guaranteed a 700ms window free of interference around each tone analyzed. After the mice became satiated and stopped engaging in the task around 30-40 minutes after the start of the experiment, pure tones of different frequencies (4, 7, 11, 17 and 23kHz) were played to the mice with each tone being 0.5-1.5 seconds apart from each other. Passively presented tones provided baseline neural responses to sounds when no movement and expectation were involved.

### Electrophysiology data collection and analysis

For electrophysiological recordings, mice were positioned in the behavioral apparatus and a 128-channel electrode (128AxN sharp, Masmanidis Lab) was lowered into the auditory cortex orthogonal to the pial surface. The electrode was connected to a digitizing head stage (Intan) and electrode signals were acquired at 30kHz, monitored in real time, and stored for offline analysis (OpenEphys). The probe was allowed to settle for 10-15 minutes and the recording started afterwards.

After recording, electrical signals were processed and the action-potentials of individual neurons were sorted using Kilosort 2.5 ^41^ and manually reviewed in Phy2 based on reported contamination, waveform principal component analysis, and inter-spike interval histograms. Tone-evoked action potential responses were measured in 1ms bins aligned to sound onset for each neuron, for each tone type, and for movement and rest conditions. Units that showed a significant (p<.01) response strength, calculated as the increase in firing rate from baseline (average over 50ms prior to stimulus) during the sound response window (0-50ms post-stimulus), to a certain frequency during passive listening were determined as sound responsive to that tone. To measure the modulation of each neuron’s tone responses to self-generated tones, we compared the neural sound response in our analysis window to the same sound in the active and passive conditions using a radial modulation index. Radial modulation was calculated as the theta value resulting from a cartesian to polar transformation of the response strength in the active condition compared to the response strength in the passive condition. Theta values were converted to a scale of +/- 2 and rotated such that a value of 0 corresponded to equal responses across the two conditions. For most analyses, only units considered responsive to a frequency were used for calculating and comparing the tone modulation index for that frequency. This inclusion criterion allowed us to focus on tone-representing units and avoid large outlier values calculated from units with firing rates always close to zero. For analysis in Fig. 2 and Fig. 3, units were chosen according to either their movement modulation group or cluster identification disregarding their sound responsiveness.

To measure the movement-related activity of each unit, we calculated the firing rate change between a 50ms window right before the onset of self-generated tones and the window far from it (450-500ms before the onset of self-generated tones). All toned trials were included regardless of the frequency of tone heard on that trial. Units that showed a significant (p<.01) increase in baseline firing rate between these two windows across trials were considered “positively modulated by movement”; neurons that showed a significant decrease were “negatively modulated”; and neurons that showed no significant change were “not modulated”. We further measured a movement modulation index in a similar way to that used for tone modulation index (see above). In practice, using either raw changes in firing rate or movement modulation index gives similar results for all analyses.

We sorted neurons into regular-spiking (RS) and fast-spiking (FS) based on measurements taken from their action potential waveforms. In particular, the duration between the trough and peak of a spike was used to determine if a unit was RS (duration>0.4ms) or FS (duration <0.4ms).

The local field potential (LFP) was extracted from recorded electrophysiological data using a low-pass filter (Equiripple, frequency limit = 200Hz). LFP signals from channels of the same depth were averaged. The tone-evoked response in LFP was measured by aligning to the onset of tone of 7kHz tones presented passively. Inverse current source density (iCSD) was calculated by taking the second derivative of the tone-evoked LFP. We assigned layers (layer1 or above, layer2/3, layer4, layer5, layer6 or lower) to different channels according to the first sink and source seen after tone onset (see Audette et al., 2022 for more detail) and each unit was considered to be at the layer of the electrode channel where it showed the largest signal.

### Non-negative factorization (NMF) and k-Means clustering

We combined non-negative factorization (NMF) and k-Means clustering to identify functional neural groups in auditory cortex. Only regular-spiking units that have an overall firing rate more than 0.2Hz were used for this analysis. For each unit included, its averaged response for expected and unexpected frequency during passive listening (500ms before and 500ms after the onset of tone, 1ms bin) was first concatenated and normalized by dividing the maximum value. This then led us to a 930-neuron-by-2000-time-bin matrix. The matrix was factorized with non-negative basis and weights using MATLAB (Statistics and Machine Learning Toolbox). To decide which rank is the best for our dataset, we ran NMF repeatedly 10 times for each rank from 0 to 25. The remaining error resulting from the factorization decreased with increasing rank used, with rank larger than 11 showing less decrease (Supp. Fig. 4A). We also compared the similarity between the resulting basis from different runs for each rank by measuring the percentage of highly correlated basis vectors between runs (Supp. Fig. 4B). We reasoned that if the rank used for NMF is higher than the true dimensions of the dataset, one or more dimensions would be forced to be further factorized to meet the rank requirement, leading to intractable results between different runs. The stability between repetitions of a specific rank thus provided us the confidence of the factorization to capture the latent variable of the dataset. We used rank=14 for our final analysis, which was the largest rank (the most variance captured) to still have a highly similar basis with different repetitions. The coordinates of units in the NMF space, which is now a neuron-by-basis matrix, were then used for k-Means clustering with MATLAB. Similarly, to determine the best number of clusters for the data, we ran k-Means repeatedly 10 times for 8-16 clusters. We measured the mutual information between clustering results across repetitions for each cluster number and chose 12 clusters, which had the most mutual information shared between runs, in our final analysis (Supp. Fig. 4C). To further make sure that we identify the best cluster for the units, we also chose the repetition that shared the most similarity in cluster assignment among the 10 repetitions. As a final move to assure units were clustered correctly, we compared the distance of units to the centroids of their assigned cluster to that to a random centroid (Supp. Fig. 4D). Units that were far away from their clusters’ centroids were then excluded from the analysis. This resulted in a final dataset of 822 units.

We also assigned manually to each cluster a cluster Type according to its response to expected and unexpected tone. The two clusters that responded rapidly and very transiently to both expected and unexpected frequencies were considered Type1. The five clusters that had rapid and rather prolonged responses compared to Type1 were considered Type2. Further, within Type2, the three clusters that showed stronger responses (higher peak) to expected tone were considered Type2Ex, while the other two showed stronger responses to unexpected tone were Type2Un. The two clusters that showed late response peaks to both frequencies were considered Type3. The three clusters that showed negative responses were considered Type4.

To measure if a cluster or cluster type showed predictive movement-related change or frequency-specific suppression, we compared the raw movement-related baseline firing rate change as well as tone modulation index of all units belonging to that group between pre- and post-learning recordings. The predictive movement-related change of Type2 clusters and tone response of Type1 clusters were then used to learn if expectation- and error-like signals were correlated. For each recording, mean movement-related baseline firing rate changes of units belonging to Type2Ex or Type2Un group were calculated for each trial with expected tone and the difference of movement-related change between the two groups was used as the measure of expectation strength. For fig3b, trials from the post-learning recording of the example mouse were sorted by their expectation strength. Mean responses of all Type1 units of the 10 trials with the weakest or strongest expectation were then calculated, soothed with a 5ms window, and shown in the figure. For fig3c, tone suppression was measured as the firing rate in a 50ms window after expectation tone onset, with lower firing representing more suppression received for that trial. Pearson correlation was calculated between movement-related change and tone suppression and compared within each mouse between pre- and post-learning recordings.

### Silencing of auditory cortex using Muscimol

Muscimol (Sigma-Aldrich) was first dissolved in distilled water to 50 mg/ml stocking solution, stored at -20oC and used within 1 month. On the same day of use, Muscimol stocking solution was diluted with saline into 0.5mg/ml working solution and kept at room temperature for at least 1 hour before application. Before the starting of each sound training session of mice used for auditory cortex silencing experiments, Muscimol was applied to auditory cortex using a little piece of Gelfoam (Pfizer) soaked with Muscimol working solution placed on the surface of the brain through the same craniotomy used for electrophysiology recording. A 15-minute waiting time was forced between the application and the beginning of the task to allow effective silencing of auditory cortex. During training, Gelfoam was kept on the brain to keep auditory cortex inactive throughout the session. After training, Gelfoam was removed and the surface of the brain was gently washed using a drop of saline. The cranial window was then sealed with a silicone elastomer (Kwik-Sil).

### Fictive pairing of M2 activation and tone

During pairing, mice were head-fixed and not required to perform a task. M2 was activated through the implanted fiber using a 470nm LED (Thorlabs, M470F3) and a 400um optic fiber cord (Thorlabs, FT400UMT). Light pulses of 15ms duration were delivered at a rate of 40Hz and power of 5mW for a total of 250ms. A 50ms tone at 7kHz was played with a 100ms delay to the starting of light stimulation. Light-tone pairings were delivered at an average rate of 0.1Hz. All mice were trained for 14-15 sessions over a week and within each session the mice experienced about 750 repetitions of light-tone pairings. During recording, mice first experienced 50 repetitions where light activation was paired with a7kHz tone. Then tones of all frequencies (4, 7, 11, 17, 23 and 41kHz) were played randomly with M2 activation for at least 50 repetitions for each frequency. Afterwards, tones of all frequencies were played to mice randomly without light stimulation. At the end of the recording, we delivered light stimulation alone without any tone to capture the response of auditory cortex following M2 activation. Neural activities were recorded and analyzed in the same way as for lever-trained mice.

### Principal component analysis (PCA) of neural dynamics

Pre- and post-learning recordings were considered as two independent datasets but were analyzed in the same way. To better focus on dimensions that distinguish between movement state and frequencies, we used only the data within a 200ms window around the tone onset (100ms before and 100ms after) for defining the PCA space. For pre-processing, each unit’s averaged response to expected and unexpected tone during movement and passive listening were downsampled to 50Hz, root-squared, smoothed with a 50ms Gaussian window and concatenated as a 1-by-40 vector. The pre-processed data were compiled and the resulting matrix was factored using MATLAB. For both pre- and post-learning data, the first 3 principal components (PCs) explained more than 90% of the variance. We then projected the full length of raw data (500ms before and after tone onset, 1ms bin) to PCA space to check how each PC contributed to discriminating movement and frequencies (Fig6b) and plotted the neural trajectories of the four conditions on the plane formed by PC2 and PC3 (Fig6a, but only showing the trajectories of 300ms before and 150ms after tone onset).

### Alignment of movement and frequency dimensions

We defined the movement dimension at a time point as the vector that points from population firing pattern during passive listening to the population firing pattern during lever press in a corresponding time bin (e.g. 20ms before tone onset) in the high dimensional neural-temporal space. Since the baseline firing pattern during rest remained relatively stable, the movement dimension always follows the trajectory of movement-related activity. We defined the frequency dimension as a similar vector to the movement dimension, but originated from the firing in passive listening to unexpected tone and points to that in passive listening to expected tone. In practice, movement dimension was only calculated for time bins prior to tone onset while frequency dimension was only for the time window 20ms to 80ms after tone onset, which was chosen to have the most discriminability between expected and unexpected frequencies according to PCA analysis. Units with an average firing rate lower than 0.2Hz were excluded from the analysis. For each unit used, its response (500ms before and after tone onset, 1ms bin) to expected and unexpected tone during passive listening and movement was first downsampled to 50Hz. The movement and frequency dimension were then computed accordingly and the angle between a total of 25 movement dimensions for a 500ms window before tone onset and the three frequency dimensions were measured. We averaged the angles between the movement dimensions at a time point to all three frequency dimensions to have the final alignment. According to the definition of movement and frequency dimensions, an alignment of around 90 degrees reflects the independence of the representation of movement and frequency, and an alignment of less than 90 degrees suggests movement-related activity skewed the neural dynamics in auditory cortex towards the representation of expected tone. Data from pre- and post-learning recordings, as well as from post-learning recordings of mice with auditory silencing during sound training, were each treated as a different dataset and independently analyzed with the same method.

### Quantification and statistical analysis

Throughout, animal values are denoted by a capital N while cell values are denoted by a lowercase n. For all figures, ∗ = p<0.05, ∗∗ = p < 0.01; ∗∗∗ = p<.001. One-sided, paired, Wilcoxon signed-rank tests were used for deciding tone responsiveness and two-sided, paired, Wilcoxon signed-rank tests were used for movement responsiveness. Unless otherwise reported, for all analyses comparing pre- and post-recordings, two-sided non-paired t-tests or two-sided non-paired one-way ANOVA were used depending on the number of variables being compared. Violin plots in all figures were made with a custom MATLAB toolbox ^42^.

---

**Supplementary Figure 1.**
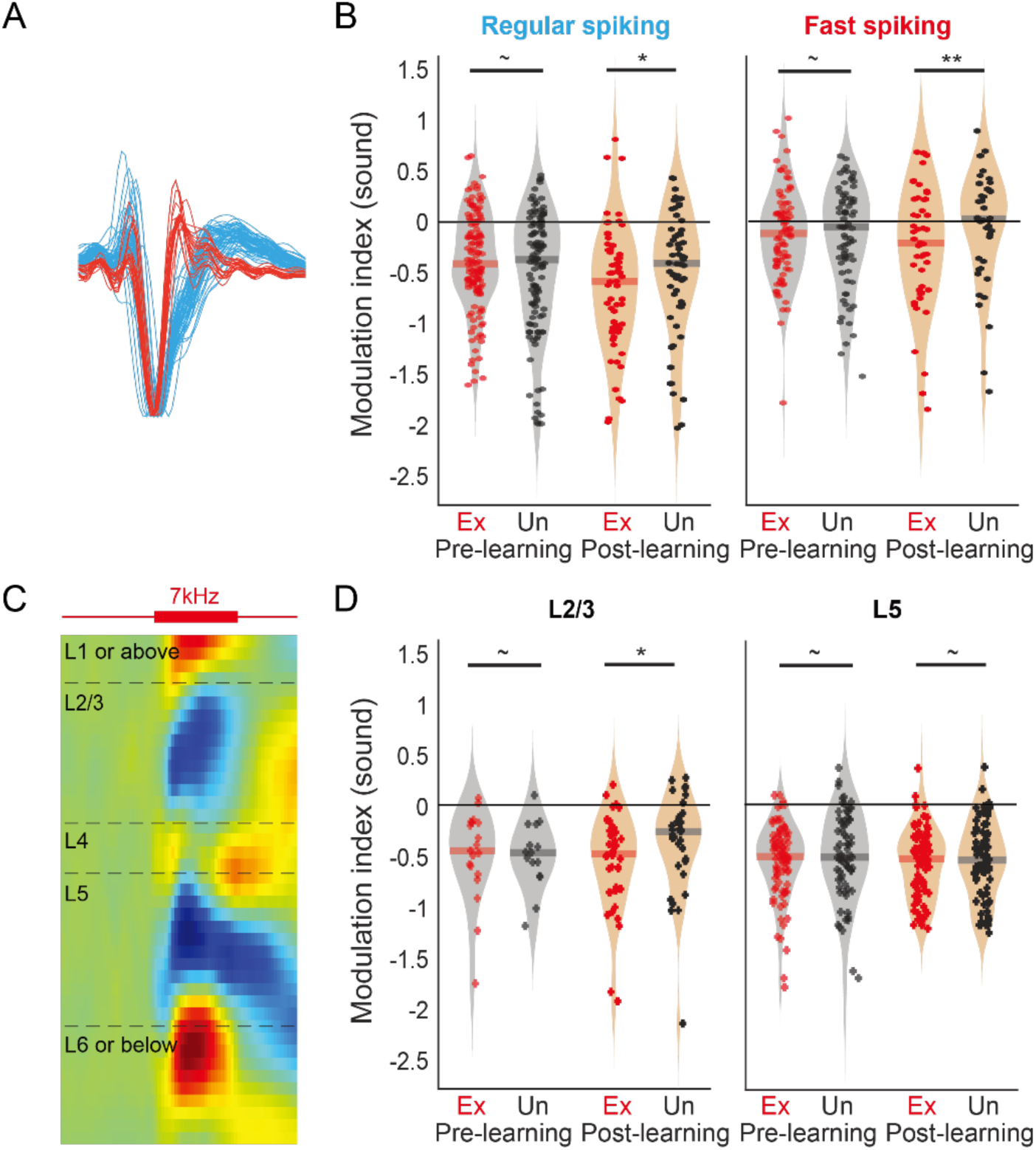
Frequency-specific suppression for different cell Types and layers. **A.** Example waveforms of regular spiking (blue) and fast spiking (red) units. **B.** Quantification of suppression to expected and unexpected tones in regular spiking or fast spiking units. Regular spiking units were more suppressed overall. Both cell types showed a frequency-specific suppression towards expected tone in post-learning sessions. **C.** Example iCSD. Different layers were identified according to the source and sink right after tone onset. **D.** Quantification of suppression to expected and unexpected tones in different layers. Frequency-specific suppression was only observed in post-learning sessions in layer2/3.

**Supplementary Figure 2.**
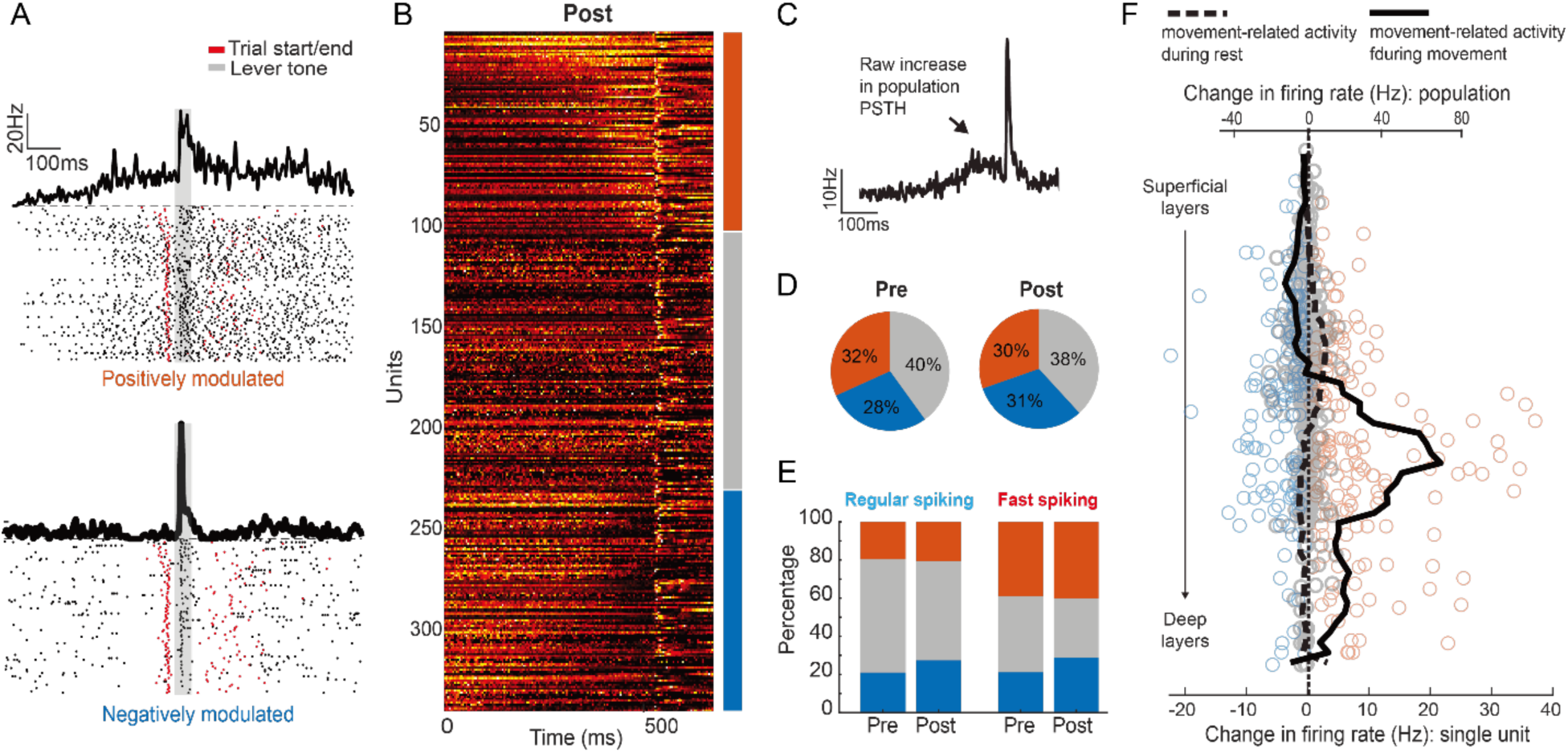
Movement-related activity in auditory cortex. **A.** Example raster plots show units that were positively or negatively modulated by movement. Change of baseline firing started prior to tone (gray shades) and trial initiation (red). **B.** Heatmap of all units recorded in post-learning sessions, sorted by the sign and starting time of their movement-related activity. A large portion of the units were affected by movement, with similar portions of units increased, decreased or kept their baseline firing prior or around the movement onset. **C.** Averaged PSTH of all units in B. Movement modulation led to an overall increase in population activity in auditory cortex. **D**. Roughly ⅓ of units were positively modulated by movement, ⅓ were negatively modulated by movement, and ⅓ unmodulated. The proportion of units showing movement-related activity were similar for pre-learning and post-learning sessions. **E.** Both regular spiking and fast spiking units showed movement-related activity, with larger portions of fast spiking units being positively modulated by movement. The portions of units affected by movement were similar for pre-learning and post-learning sessions for both cell types. **F.** Movement-related activity of each unit in B by their depth in auditory cortex. Units positively (orange circle), negatively (blue circle) or not modulated (gray circle) by movement were found across all depths, with positively modulated units more concentrated in deeper layers. During passive listening before tone onset (black dotted line), the baseline activity of auditory cortical population remained stable. In contrast, during sound-generating lever task (black line) population activity increased in deeper layers.

**Supplementary Figure 3.**
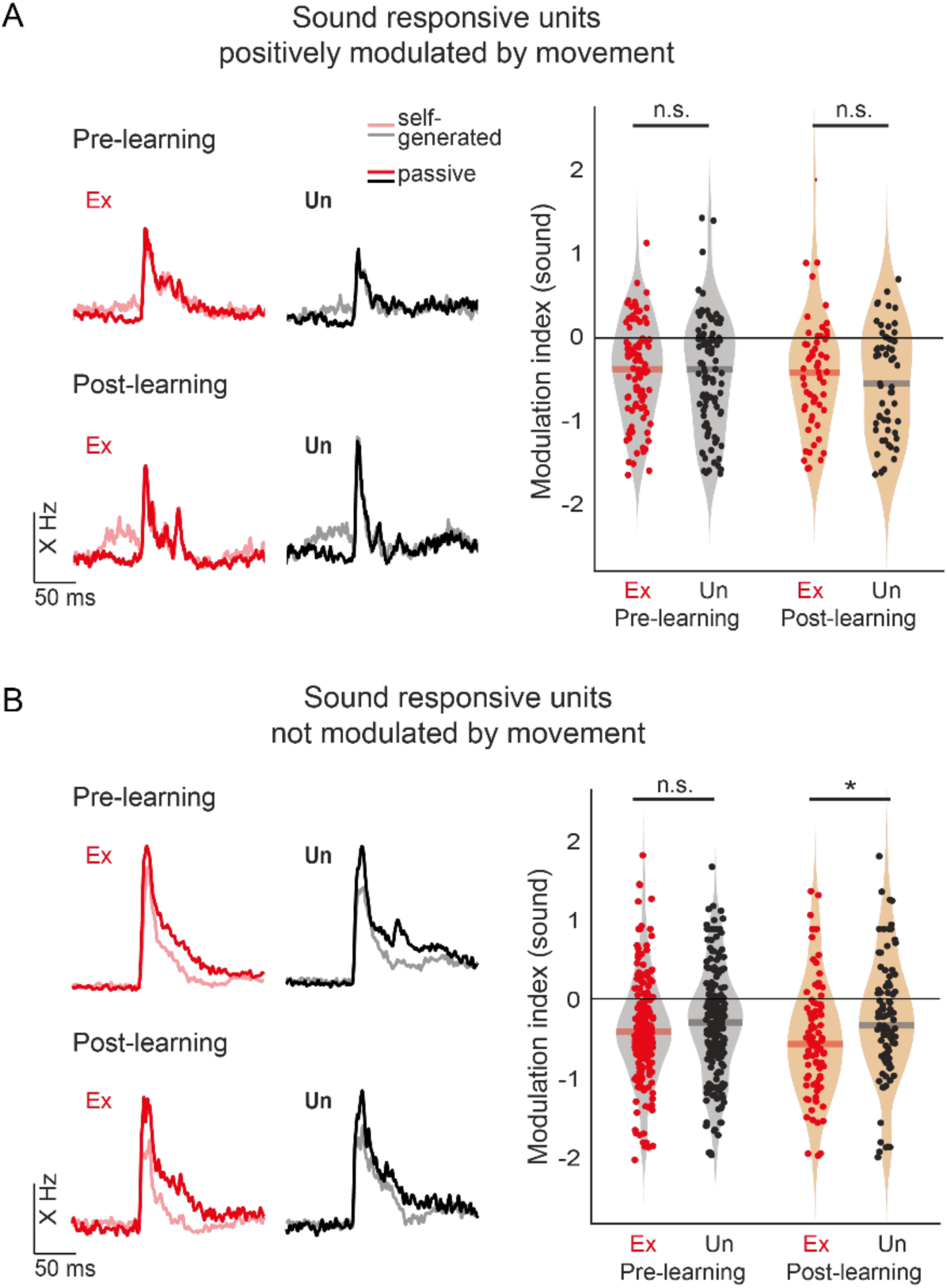
Frequency-specific suppression and movement-related expectation were observed in different neural groups. **A.** Sound responsive units that were positively modulated by movement with their population PSTH (left) and sound modulation index (right). Frequency-specific suppression was not found in this population (right, modulation index of sound). **B.** Sound responsive units that were not modulated by movement with their population PSTH (left) and sound modulation index (right). Frequency-specific suppression was found in post-learning sessions, with expected tone more strongly suppressed. This suggested a separation of movement-related expectation and frequency-specific suppression into different populations.

**Supplementary Figure 4.**
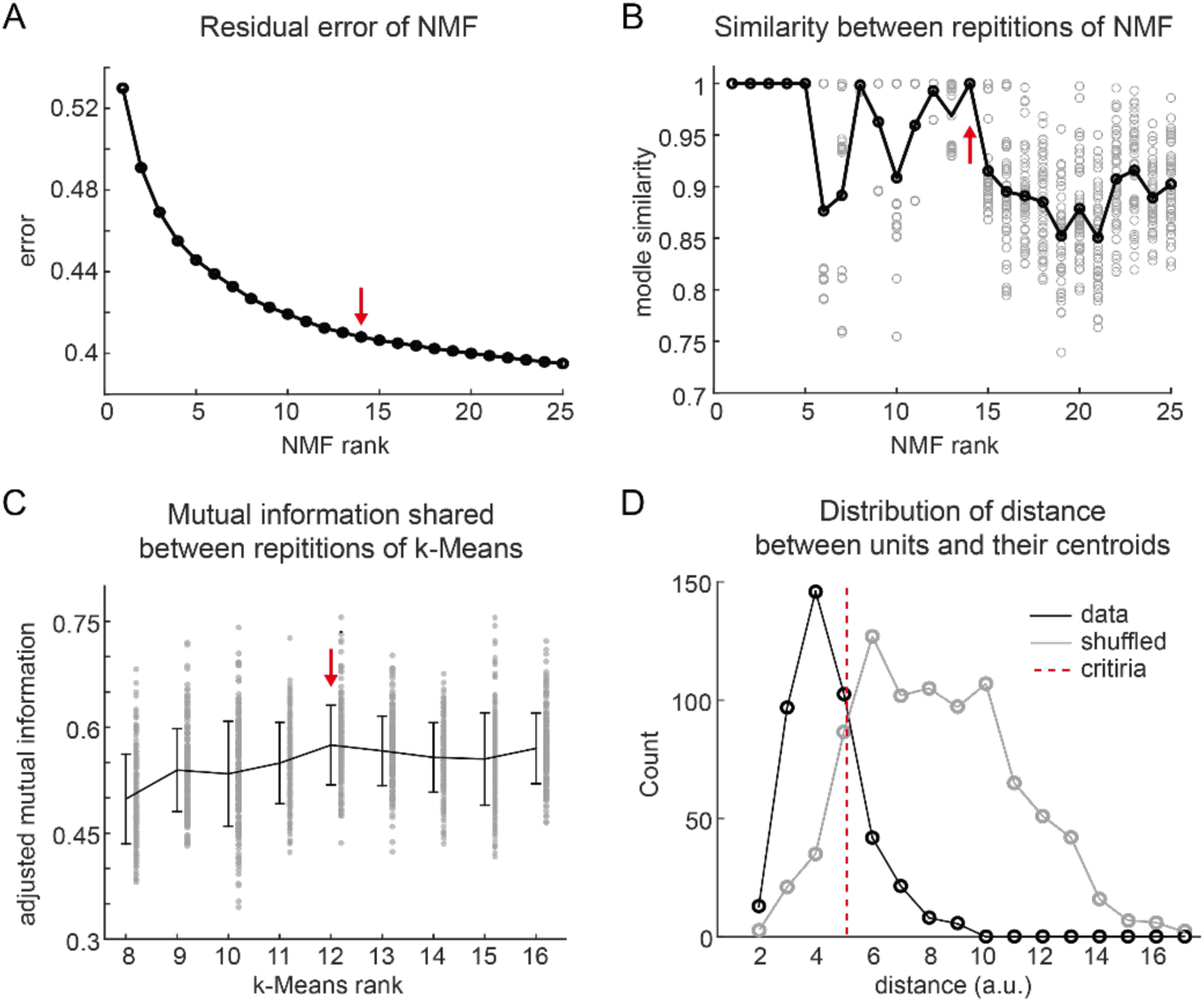
Methods for NMF-kMeans clustering. **A.** Residual error of NMF for different ranks. Red error points to the error for rank (14) used for NMF-kMeans clustering. **B.** Similarity of NMF basis between reputations of NMF analysis with the same rank. NMF results were highly stable for lower ranks but became less tractable between repetitions as the rank increases. Red error points to the similarity of rank (14) used for NMF-kMeans clustering. **C.** Mutual information shared by clustering results of different repetitions of kMeans with the same number of clusters targeted. Red error points to cluster number (12) with the highest mutual information shared, which was used for NMF-kMeans clustering. **D.** Distribution of distance of units to the centroid of their cluster assigned (black) or centroid of a random cluster (gray). A distance threshold (dotted red line) was adopted to best separate the two distributions. Clusters with distance to their clusters’ centroid that were further than the threshold were excluded from the clustering results and later analysis.

**Supplementary Figure 5.**
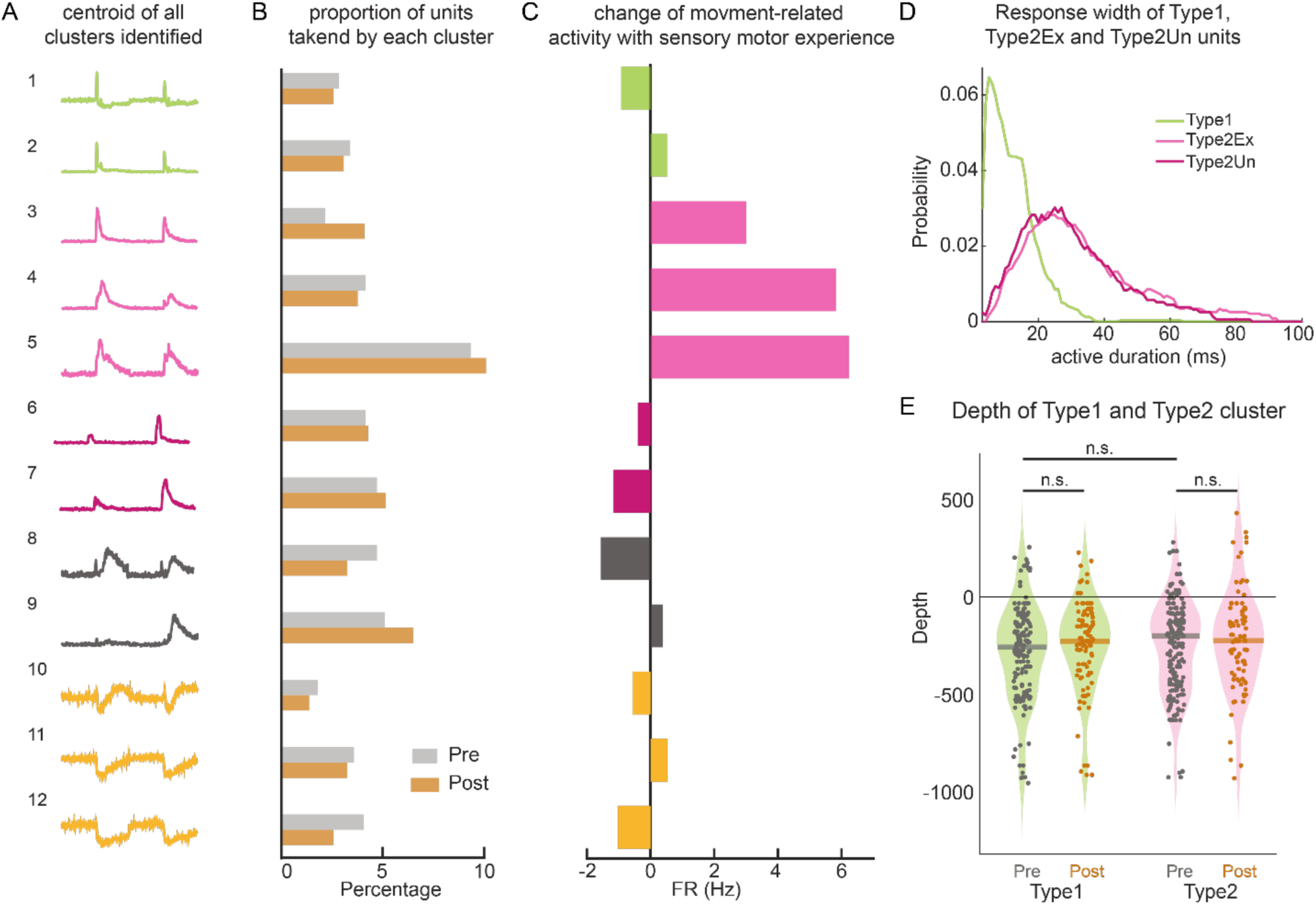
Properties of different clusters identified by NMF-kMeans clustering. **A.** Centroid of all 12 clusters from NMF-kMeans clustering. Color depicts the type of the cluster (Type1, green; Type2Ex, light pink; Type2Un, dark pink; Type3, dark gray; Type4, yellow). **B.** Percentage of units assigned to each cluster were similar for pre-learning and post-learning sessions. **C.** Difference in movement-related activity between pre-learning and post-learning sessions. Most clusters showed similar movement-related activity for both sessions, except for the three clusters of Type2Ex, which showed a great increase in movement-related activity in post-learning sessions. **D.** Response width of Type1 (green), Type2Ex (light pink) and Type2Un (dark pink) units. Both Type2Ex and Type2Un units showed longer duration of activation by sound compared to Type1 units. **E.** Type1 and Type2 units were found across layers and showed similar distribution of depth between the two types as well as between pre- and post-learning sessions.

**Supplementary Figure 6.**
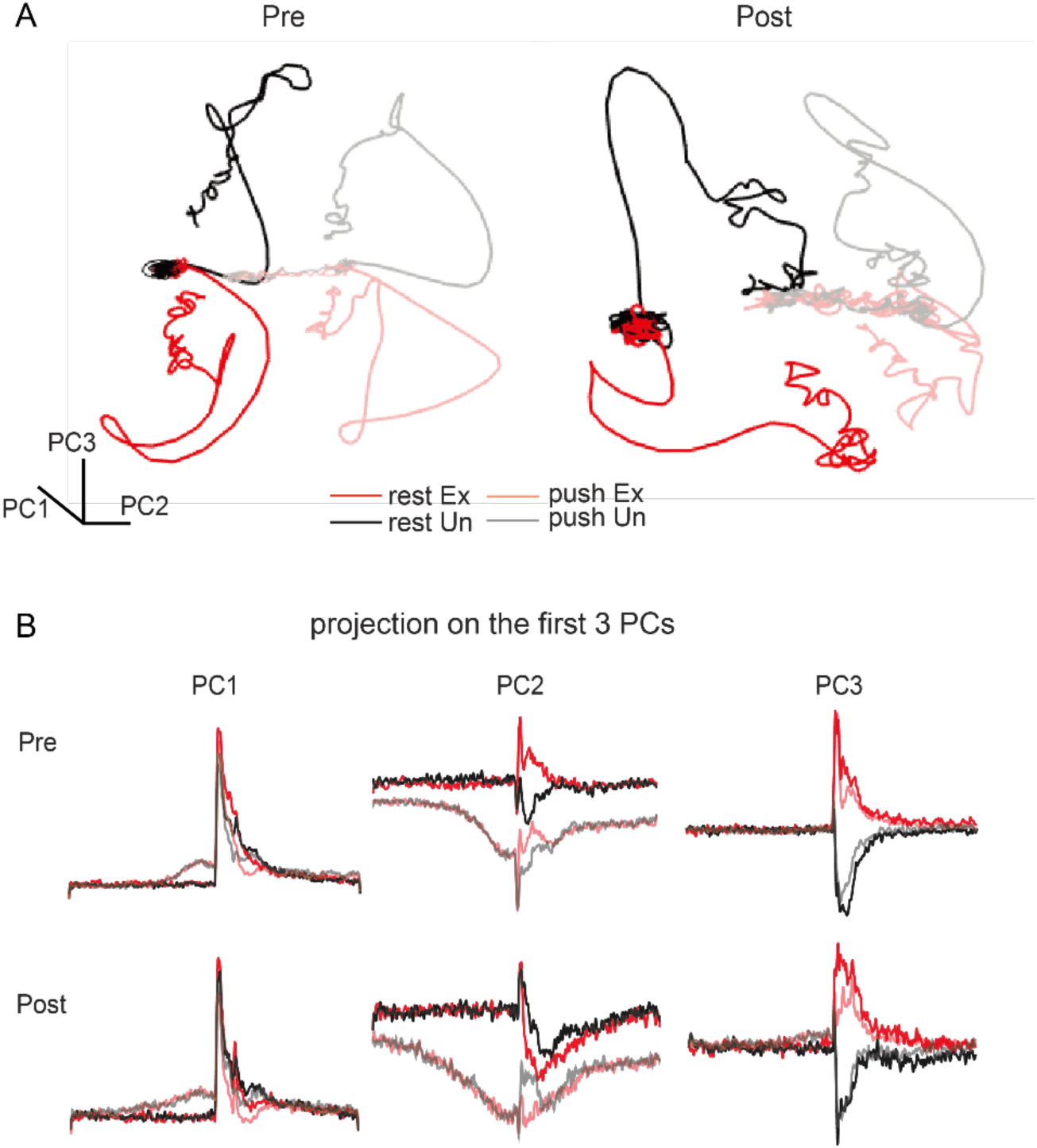
Full projections of the PCA analysis. **A.** Neural trajectories of responses to expected (red) and unexpected (black) tones during rest and during movement (semi-transparent) in pre-learning (left) and post-learning (right) sessions, projected to the space formed by the first 3 PCs. **B.** Projections of pre-learning and post-learning data to each of the first 3 PCs. Activity on PC1 showed large tone-evoked responses indifferent to frequencies. Activity on PC2 were of opposite sign for responses during rest and movement, with a large movement-related change prior to tone onset for response during movement. Activity on PC3 were of opposite sign for expected and unexpected frequencies.

